# Genome-wide association study in the pseudocereal quinoa reveals selection pattern typical for crops with a short breeding history

**DOI:** 10.1101/2020.12.03.410050

**Authors:** Dilan S. R. Patiranage, Elodie Rey, Nazgol Emrani, Gordon Wellman, Karl Schmid, Sandra M. Schmöckel, Mark Tester, Christian Jung

## Abstract

Quinoa germplasm preserves useful and substantial genetic variation, yet it remains untapped due to a lack of implementation of modern breeding tools. We have integrated field and sequence data to characterize a large diversity panel of quinoa. Whole-genome sequencing of 310 accessions revealed 2.9 million polymorphic high confidence SNP loci. Highland and Lowland quinoa were clustered into two main groups, with *F*_*ST*_ divergence of 0.36 and fast LD decay of 6.5 and 49.8 Kb, respectively. A genome-wide association study uncovered 600 SNPs stably associated with 17 agronomic traits. Two candidate genes are associated with thousand seed weight, and a resistance gene analog is associated with downy mildew resistance. We also identified pleiotropically acting loci for four agronomic traits that are highly responding to photoperiod hence important for the adaptation to different environments. This work demonstrates the use of re-sequencing data of an orphan crop, which is partially domesticated to rapidly identify marker-trait association and provides the underpinning elements for genomics-enabled quinoa breeding.

## Introduction

Climate change poses a great threat to crop production worldwide. In temperate climates of the world, higher temperatures and extended drought periods are expected. Moreover, crop production in industrialized countries depends on only a few major crops resulting in narrow crop rotations. Therefore, rapid transfer of wild species into crops using genetic modification and targeted mutagenesis is currently discussed ^1,2^. Alternatively, orphan crops with a long tradition of cultivation but low breeding intensity can be genetically improved by genomics assisted selection methods. Quinoa (*Chenopodium quinoa* Willd.) is a pseudocereal crop species with a long history of cultivation. It was first domesticated about 5000-7000 years ago in the Andean region. Quinoa was a staple food during the pre-Columbian era, and the cultivation declined after the introduction of crops like wheat and barley by the Spanish rulers. Owing to diversity, biotic and abiotic stress tolerance, and ecological plasticity, quinoa can adapt to a broad range of agroecological regions ^3,4^. Due to a high seed protein content and a favorable amino acid composition, its biological value is even higher than beef, fish, and other major cereals ^5,6^. These favorable characteristics contributed to the increasing worldwide popularity of quinoa among consumers and farmers.

A spontaneous hybridization event between two diploid species between 3.3 and 6.3 million years ago gave rise to the allotetraploid species quinoa (2*n* = 4*x* = 36) with a genome size of 1.45-1.5 Gb (nuclear DNA content 1C = 1.49 pg) ^7,8^. A reference genome of the coastal Chilean quinoa accession PI 614886 has been published with 44,776 predicted gene models together with whole-genome re-sequencing of *C. pallidicaule* and *C. suecicum* species, close relatives of the A and B subgenome donor species, respectively *^9^.* The organellar genomes are originated from the A-genome ancestor ^10^.

Quinoa belongs to the Amaranthaceae, together with some other economically important crops like sugar beet, red beet, spinach, and amaranth. It reproduces sexually after self-pollination. Facultative autogamy was reported for plants in close proximity with outcrossing rates in a range of 0.5 to 17.36 % ^11,12^. Thus, quinoa accessions are typically homozygous inbred lines. Nonetheless, heterozygosity in some accessions has been reported, which indicates cross-pollination ^13^. The inflorescences are panicles, which are often highly branched. Florets are tiny, which is a significant obstacle for hand-crossing. However, routine protocols for F1 seed production in combination with marker-assisted selection have been developed recently ^14,15^.

Systematic breeding of quinoa is still at its infancy compared to major crops. Until recently, breeding has been mainly limited to Bolivia ^16^ and Peru ^17^, which are the major growing areas of quinoa. Therefore, quinoa can be regarded as a partially domesticated crop. Many accessions suffer from seed shattering, branching, and non-appropriate plant height, which are typical domestication traits. Apart from these characters, grain yield and seed size, downy mildew resistance, synchronized maturity, stalk strength, and low saponin content are major breeding objectives ^18^. In the past years, activities have been intensified to breed quinoa genotypes adapted to temperate environments, for example, Europe, North America, and China ^19^. Here, the major problem is the adaptation to long-day conditions because quinoa is predominantly a short-day plant due to its origin from regions near the equator.

There are only a few studies about the genetic diversity of quinoa. They were mainly based on phenotypic observations ^16,20^ and low throughput marker systems like random amplified polymorphic DNA ^21^, amplification fragment length polymorphisms ^22^, and microsatellites ^23^. A limited number of single nucleotide polymorphisms (SNP) based on expressed sequence tags were published ^24^. Maughan, et al. ^25^ used five bi-parental populations to identify ca. 14,000 SNPs, from which 511 KASP markers were developed. Genotyping 119 quinoa accessions gave the first insight into the population structure of this species ^25^. Now, the availability of a reference genome enables genome-wide genotyping (Jarvis et al. 2017). Jarvis, et al. ^9^ re-sequenced 15 accessions and identified ca. 7.8 million SNPs. In another study, 11 quinoa accessions were re-sequenced, and 8 million SNPs and ca. 842 thousand indels were identified ^26^.

Our study aimed to analyze the population structure of quinoa and patterns of variation by re-sequencing a diversity panel encompassing germplasm from all over the world. Using millions of markers, we performed a genome-wide association study using multiple-year field data. Here, we identified QTLs that control agronomically important traits important for breeding cultivars to be grown under long-day conditions. We are discussing the fundamental differences between an underutilized crop and crops with a long breeding history. Our results provide useful information for further understanding the genetic basis of agronomically important traits in quinoa and will be instrumental for future breeding.

## Results

### Re-sequencing 310 quinoa accessions reveals high sequence variation

We assembled a diversity panel made of 310 quinoa accessions representing regions of major geographical distributions of quinoa (Supplementary Fig. 1). The diversity panel comprises accessions with different breeding history (Supplementary Table 1). We included 14 accessions from a previous study, of which 7 are wild relatives ^9^. The sequence coverage ranged from 4.07 to 14.55, with an average coverage of 7.78. We mapped sequence reads to the reference genome V2 (CoGe id53523). Using mapping reads, we identified 45,330,710 single nucleotide polymorphisms (SNPs).

After filtering the initial set of SNPs, we identified 4.5 million SNPs in total for the base SNP set. We further filtered the SNPs for MAF >5 % (HCSNPs). We obtained 2.9 million high confident SNPs for subsequent analysis (Supplementary Table 2). Across the whole genome, SNP density was high, with an average of 2.39 SNPs/kb. However, SNP densities were highly variable between genomic regions and ranged from 0 to 122 SNPs/kb (Supplementary Fig. 3). We did not observe significant differences in SNP density between the two subgenomes (A subgenome 2.43 SNPs/kb; B subgenome 2.35 SNPs/kb). Then, we split the SNPs by their functional effects as determined by SnpEff ^27^. Among SNPs located in non-coding regions, 598,383 and 617,699 SNPs were located upstream (within 5kb from the transcript start site) and downstream (within 5kb from the stop site) of a gene, whereas 114,654 and 251,481 SNPs were located within exon and intron sequences, respectively (Table 1). We further searched for SNPs within coding regions. We found 70,604 missense SNPs and 41,914 synonymous SNPs within coding regions of 53,042 predicted gene models.

**Table 1:** Summary statistics of genome-wide single nucleotide polymorphisms identified in 303 quinoa accessions

| Parameter | Type | All genotypes<br>(quinoa only) | Highland<br>population | Lowland<br>population |
| --- | --- | --- | --- | --- |
| SNP | Total | 2,872,935 | 2,590,907 | 1,938,225 |
|  | Intergenic | 2,452,347 | 2,227,952 | 1,649,310 |
|  | Introns | 251,481 | 101,546 | 172,692 |
|  | Exons | 114,654 | 214,945 | 78,248 |
| Nucleotide diversity | | | $5.78 \times 10^{-4}$ | $3.56 \times 10^{-4}$ |
| Tajima's *D* |  |  | 0.884 | -0.384 |
| Population<br>divergences | *F<sub>ST</sub>*<br>(Weighted average) |  | 0.36 |  |

### Linkage disequilibrium and population structure of the quinoa diversity panel

Across the whole genome, LD decay between SNPs averaged 32.4 kb. We did not observe substantial LD differences between subgenome A (31.9kb) and subgenome B (30.7kb) (Supplementary Fig. 4C). The magnitude of LD decay among chromosomes did not vary drastically except for chromosome Cq6B, which exhibited a substantially slower LD decay (Supplementary Fig. 4 A and B).

Then, we unraveled the population structure of the diversity panel. We performed principal component (PCA(SNP)), population structure, and phylogenetic analyses. PCA(SNP) showed two main clusters consistent with previous studies ^13^. The first and second principal components (PC1(SNP) and PC2(SNP)) explained 23.35% and 9.45% of the variation, respectively (Fig. 1A). 202 (66.67%) accessions were assigned to subpopulation 1 (SP1) and 101 (33.33%) to subpopulation 2 (SP2). SP1 comprised mostly Highland accessions, whereas Lowland accessions were found in SP2. PCA demonstrated a higher genetic diversity of the Highland population (Fig. 1A). We also calculated PCs for each chromosome separately. For 16 chromosomes, the same clustering as for the whole genome was calculated. Nevertheless, two chromosomes, Cq6B, and Cq8B showed three distinct clusters (Supplementary Fig. 5). This is due to the split of the Lowland population into two clusters. We reason that gene introgressions on these two chromosomes from another interfertile group might have caused these differences. This is also supported by a slower LD decay on chromosome Cq6B (Supplementary Fig. 4B).

**Fig. 1:**
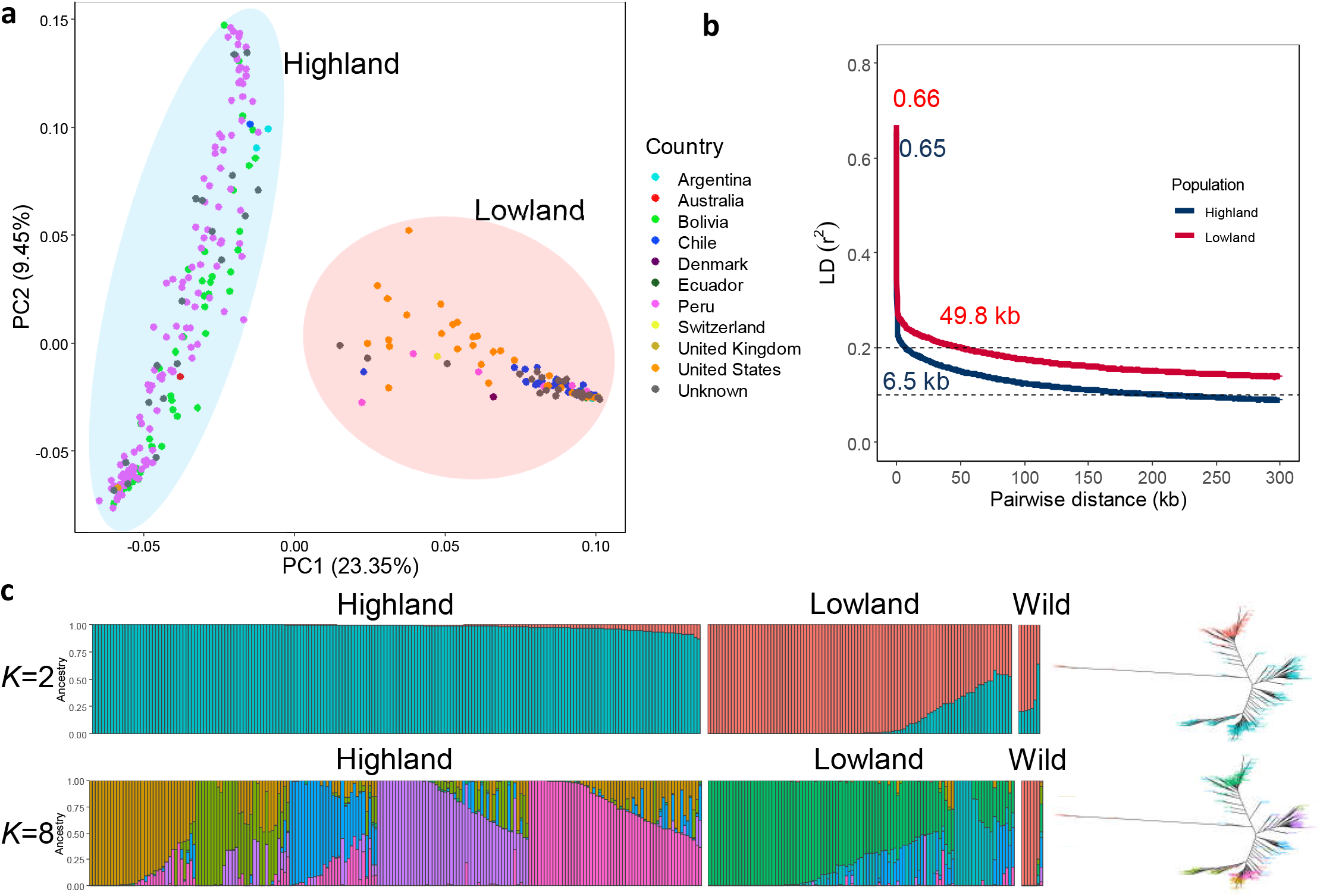
Genetic diversity and population structure of the quinoa diversity panel. (a) PCA of 303 quinoa accessions. PC1 and PC2 represent the first two components of analysis, accounting for 23.35% and 9.45% of the total variation, respectively. The colors of dots represent the origin of accessions. Two populations are highlighted by different colors: Highland (light blue) and Lowland (pink). (b) Subpopulation wise LD decay in Highland (blue) and Lowland population (red). (c) Population structure is based on ten subsets of SNPs, each containing 50,000 SNPs from the whole-genome SNP data. Model-based clustering was done in ADMIXTURE with different numbers of ancestral kinships (*K*=2 and *K*=8). *K*=8 was identified as the optimum number of populations. Left: Each vertical bar represents an accession, and color proportions on the bar correspond to the genetic ancestry. Right: Unrooted phylogenetic tree of the diversity panel. Colors correspond to the subpopulation.

We also performed a population structure analysis with the ADMIXTURE software. We used cross-validation to estimate the most suitable number of populations. Cross-validation error decreased as the *K* value increased, and we observed that after *K* = 5, cross-validation error reached a plateau (Supplementary Fig. 6B). We observed allelic admixtures in some accessions, likely owing to their breeding history. The wild accessions were also clearly separated at the smallest cross-validation error of *K*=8, except two *C. hircinum* accessions (Fig. 1C). The reason for this could be that because *C. hircinum* is the closest crop wild relative, it also may have outcrossed with quinoa. The Highland population was structured into five groups, while the Lowland accessions were split into two subpopulations. The broad agro-climatic diversity of the Andean Highland germplasm might have caused a higher number of subpopulations.

We analyzed the phylogenetic relationships between quinoa accessions using 434,077 SNPs. Constructing a maximum likelihood tree gave rise to five clades (Fig. 2). We found that the placement of the wild quinoa species was concordant with the previous reports confirming that quinoa was domesticated from *C. hircinum* ^9^. However, we found that the *C. hircinum* accession BYU 566 (from Chile) was placed at the base of both Lowland and Highland clades, which is in contrast to Jarvis, et al. ^9^, where this accession was placed at the base of coastal quinoa. As expected, accessions from the USA and Chile are closely related because the USDA germplasm had been collected at these geographical regions.

**Fig. 2:**
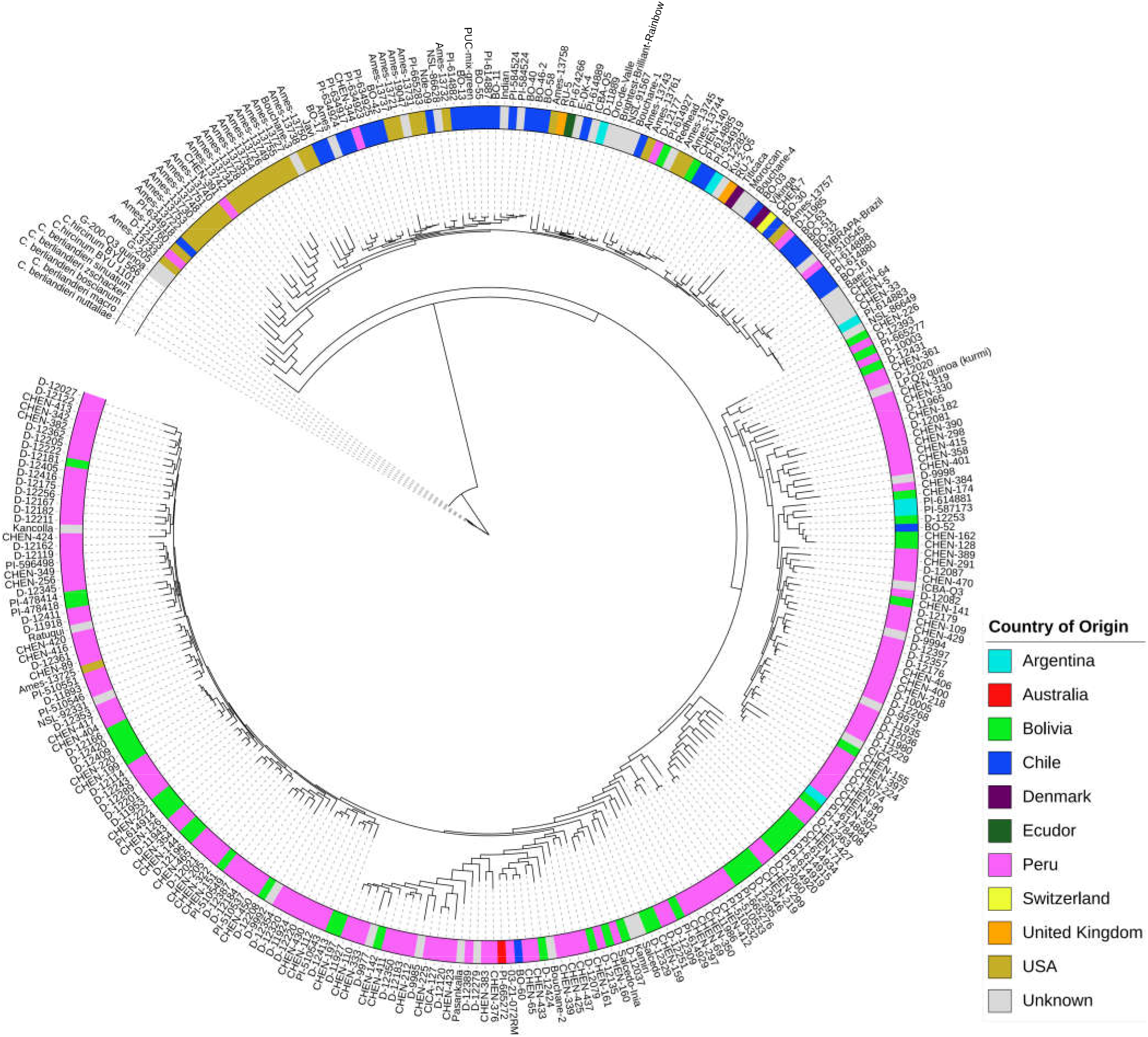
Maximum likelihood tree of 303 quinoa and seven wild *Chenopodium* accessions from the diversity panel. Colors are depicting the geographical origin of accessions.

### Genomic patterns of variations between Highland and Lowland quinoa

We were interested in patterns of variation in response to geographical diversification. We used principal component analysis derived clusters and phylogenetic analysis to define two diverged quinoa populations (namely Highland and Lowland). These divergent groups are highly correlated with Highland and Lowland geographical origin. We used the base SNP set to analyze diversity statistics. To detect genomic regions affected by the population differentiation, we measured the level of nucleotide diversity using 10 kb non-overlapping windows ^28^. Then we calculated the whole genome-wide LD decay across the two populations (Highland vs. Lowland); LD decayed more rapidly in Highland quinoa (6.5 kb vs. 49.8 kb) (Fig. 1B). To measure nucleotide diversity, we scanned the quinoa genome with non-overlapping windows of 10 kb in length in both populations separately. The nucleotide diversity of the Highland population (5.78 × 10^−4^) was 1.62 fold higher compared to the Lowland population (3.56 × 10^−4^) (Table 1 and Supplementary Fig. 7). We observed left-skewed distribution and negative Tajima’s *D* value (−0.3883) in the Lowland populations indicating recent population growth (Table 1 and Supplementary Fig. 8). Genomic regions favorable for adaptation to Highlands should have substantially lower diversity in the Highland population than the Lowland population. Therefore, we calculated the nucleotide diversity ratios between Highland and Lowland to identify major genomic regions that are underlying the population differentiation. The *F*_*ST*_ value between populations was estimated to be 0.36, illustrating strong population differentiation. Concerning the regions of variants, the number of exonic SNPs is substantially higher in the Highland population (Table 1 and Supplementary Fig. 7).

### Mapping agronomically important trait loci in the quinoa genome

We evaluated 13 qualitative and four dichotomous traits on 350 accessions across two different environments. At the time of the final harvest, 254 accessions did not reach maturity (senescence). All accessions produced seeds therefore used in seed analysis. For all traits, substantial phenotypic variation among accessions was found. High heritabilities were calculated for all quantitative traits except for number of branches (NoB) and stem lying (STL), which indicates that the phenotypic variation between the accessions is mostly caused by genetic variation (Supplementary Table 3). Trait correlations between years were also high (Supplementary Fig. 9), which is in accordance with the heritability estimates. We found the strongest positive correlation between days to maturity (DTM) and panicle length (PL), and plant height (PH) and PL, whereas the strongest negative correlation was found between DTM and thousand seed weight (TSW) (Supplementary Fig. 10). Then a principal component analysis was performed based on 12 quantitative traits (PCA_(PHEN)_) to explore the phenotypic relationship among quinoa accessions. The first two principal components explained 62.12% of the phenotypic variation between the accessions. The score plot of the principal components showed a similar clustering pattern as the SNP based PCA analysis (PCA(SNP)) (Fig. 1A and Supplementary Fig. 11A). PCA_(PHEN)_ variables factor map indicated that most Lowland accessions were high yielding with high TSW and dense panicles. Moreover, these accessions are early flowering and early maturing, and they are short (Supplementary Fig. 11B). Phenotype-based PCA_(PHEN)_ also showed that the Lowland accessions are better adapted/selected for cultivation in long-day photoperiods compared to the Highland accessions. These results are in accordance with LD, nucleotide diversity, and Tajima’s *D* estimations, implying the Lowland accessions went through a stronger selection during breeding.

Then, we calculated the best linear unbiased estimates (BLUE) of the traits investigated. In total, 294 accessions shared the re-sequencing information and phenotypes out of 350 phenotypically evaluated accessions. For GWAS analysis, we used ~2.9 million high-confidence SNPs. In total, we identified 1480 significant (*P*<9.41e-7) SNP-trait associations (MTA) for 17 traits (Supplementary Fig. 12). The number of MTAs ranged from 4 (STL) to 674 (DTM) (Supplementary Table 4). In agreement with previous reports, we defined an MTA as “consistent” when it was detected in both years^29^. We identified 600 consistent MTAs across eleven traits. TSW and DTM showed the highest number of “consistent” associations. Among these, 143 MTAs are located within a gene, and 22 SNPs resulted in a missense mutation (Supplementary Table 5). MTA for the duration from bolting to flowering (DTB to DTF), NoB, Seed yield, STL, and growth type (GT) were not “consistent” between years (Supplementary Fig. 12). This is also reflected by the low heritability estimations of these traits, indicating considerably higher genotype x environment interactions.

### Candidate genes for agronomically important traits

First, we tested the resolution of our mapping study. We searched for major genes 50Kb down-and upstream of significant SNPs for two qualitative traits in quinoa, flower color, and seed saponin content. We identified highly significant MTAs for stem color on chromosome Cq1B (69.72-69.76 Mb). There are two genes (*CqCYP76AD1* and *CqDODA1*) from the associated loci displaying high homology to betalain synthesis pathway genes *BvCYP76AD1* ^30^ and *BvDODA1* ^31^ from sugar beet (Supplementary Fig. 14A and Supplementary Fig. 12). A significant MTA for saponin content on chromosome Cq5B between 8.85 Mb to 9.2 Mb harbored the two *BHLH25* genes which have been reported to control saponin content in quinoa ^9^ (Supplementary Fig. 14B and Supplementary Fig. 12). This demonstrates that the marker density is high enough to narrow down to causative genes underlying a trait.

Then, we examined four quantitative traits. We found seven MTA on chromosome Cq2A that are associated with DTF, DTM, PH, and PL (cross-phenotype association), indicating evidence for pleiotropic gene action (Fig. 3 and Supplementary Table 6). For further confirmation and to investigate genes that are pleiotropically active on different traits, we followed a multivariate approach ^32^. First, we performed a PCA using the four phenotypes (cross-phenotypes). We found 89.94% of the variation could be explained by the first two principal components of the cross-phenotypes (PCA(CP)) (Supplementary Fig. 15). This indicates the adequate power of the PCA(CP) to reduce dimensions for the analysis of the cross-phenotypes association. We observed similar clustering as in PCA(SNP). Therefore, these results indicate that in quinoa, DTF, DTM, PH, and PL are highly associated with population structure and thus, the adaptation to diverse environments. Then, we performed a GWAS analysis using the first three PCs as traits (PC-GWAS) (Supplementary Fig. 15C). We identified strong associations on chromosomes Cq2A, Cq7B (PC1), and Cq8B (PC2) (Supplementary Fig. 16). Out of 468 MTAs (PC1:426 and PC2 42) across the whole genome, 222 (PC1:211 and PC2:11) are located within 95 annotated genes. We found 14 SNPs that changed the amino acid sequence in 12 predicted protein sequences of associated genes (Supplementary Table 5). In the next step, we searched genes located within 50kb to an MTA. Altogether, 605 genes were identified (PC1:520 and PC2:85) (Supplementary Table 7).

**Fig. 3:**
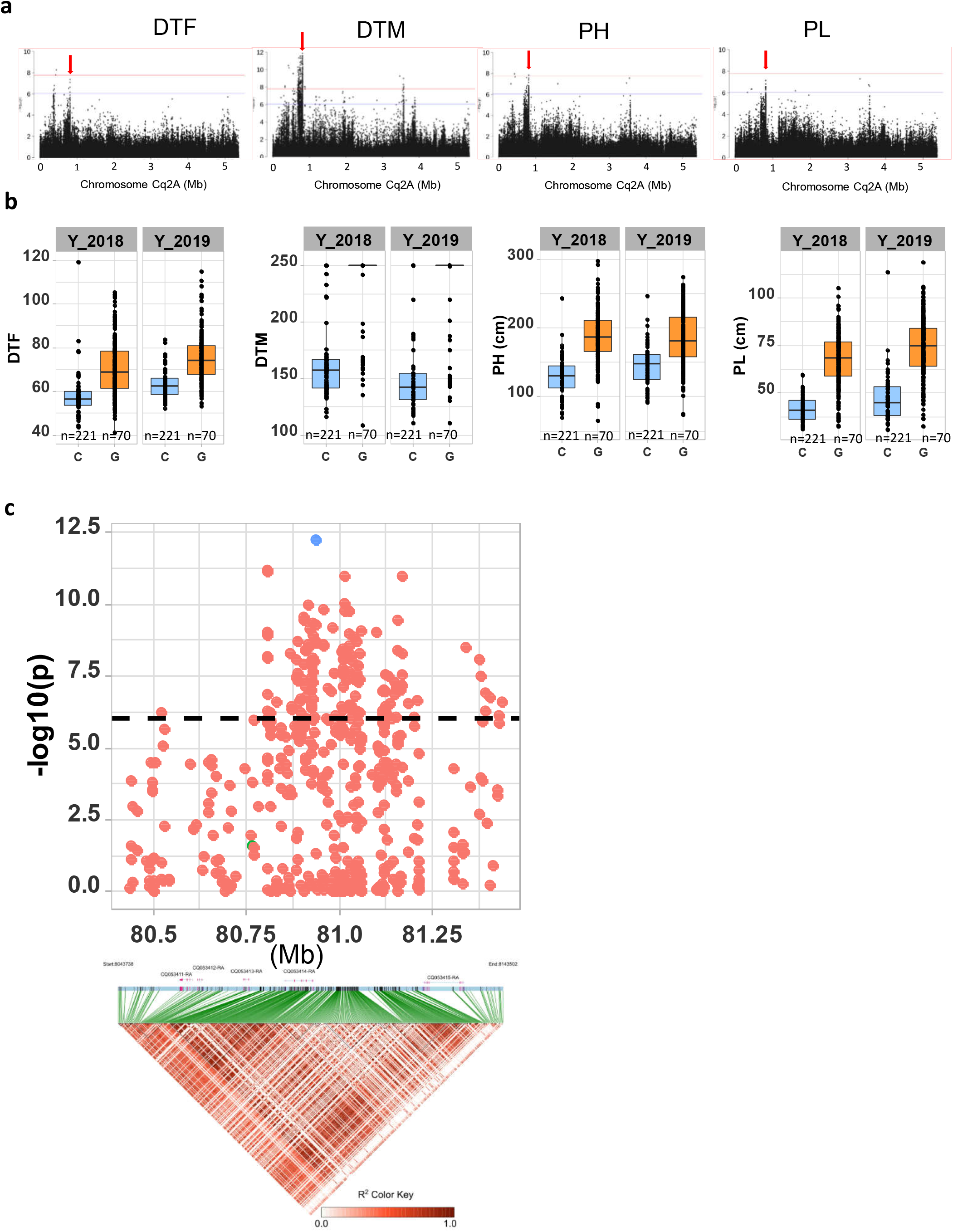
Genomic regions associated with important agronomic traits (a) Significant marker-trait associations for days to flowering, days to maturity, plant height, and panicle density on chromosome Cq2A. Red color arrows indicate the SNP loci pleiotropically acting on all four traits. (b) Boxplots showing the average performance for four traits over two years, depending on single nucleotide variation (C or G allele) within locus Cq2A_ 8093547. (c) Local Manhattan plot from region 80.40 - 81.43 Mb on chromosome Cq2A associated with PC1 of the days to flowering (DTF), days to maturity (DTM), plant height (PH), and panicle length (PL), and local LD heat map (bottom). The colors represent the pairwise correlation between individual SNPs. Green color dots represent the strongest MTA (Cq2A_ 8093547).

We found the region 80.50 −81.50 Mb on chromosome Cq2A to be of special interest because it displays stable pleiotropic MTA for DTF, DTM, PH, and PL. The most significant SNP is located within the *CqGLX2-2* gene, which encodes an enzyme of the glyoxalase family (Fig. 3). The Arabidopsis *GLX2-1* has been shown to be essential for growth under abiotic stress ^33^. The allele carrying a cytosine at the position with the most significant SNP resulted in early flowering, maturing, and short panicles and plants (Fig. 3b). These traits are essential for the adaptation to long-day conditions.

Thousand seed weight is an important yield component. We found a strong MTA between 63.2 – 64.87 Mb on chromosome Cq8B. Significantly associated SNPs were localized within two genes (Fig. 4). One gene displays homology to *PP2C* encoding a member of the phosphatase-2C (*PP2C*) protein family, which participates in Brassinosteroids signaling pathways and controls the expression of the transcription factor *BZR1* ^34^. The second gene encodes a member of the RING-type E3 ubiquitin ligase family. These genes are controlling seed size in soybean, maize, rice, soybean, and Arabidopsis ^35^. We then checked haplotype variation and identified 5 and 7 haplotypes for *CqPP2C* and *CqRING* genes, respectively. Accessions carrying PP2C_hap3 and RING_hap7 displayed larger seeds in both years (Fig. 4 and Supplementary Fig. 17)

**Fig. 4:**
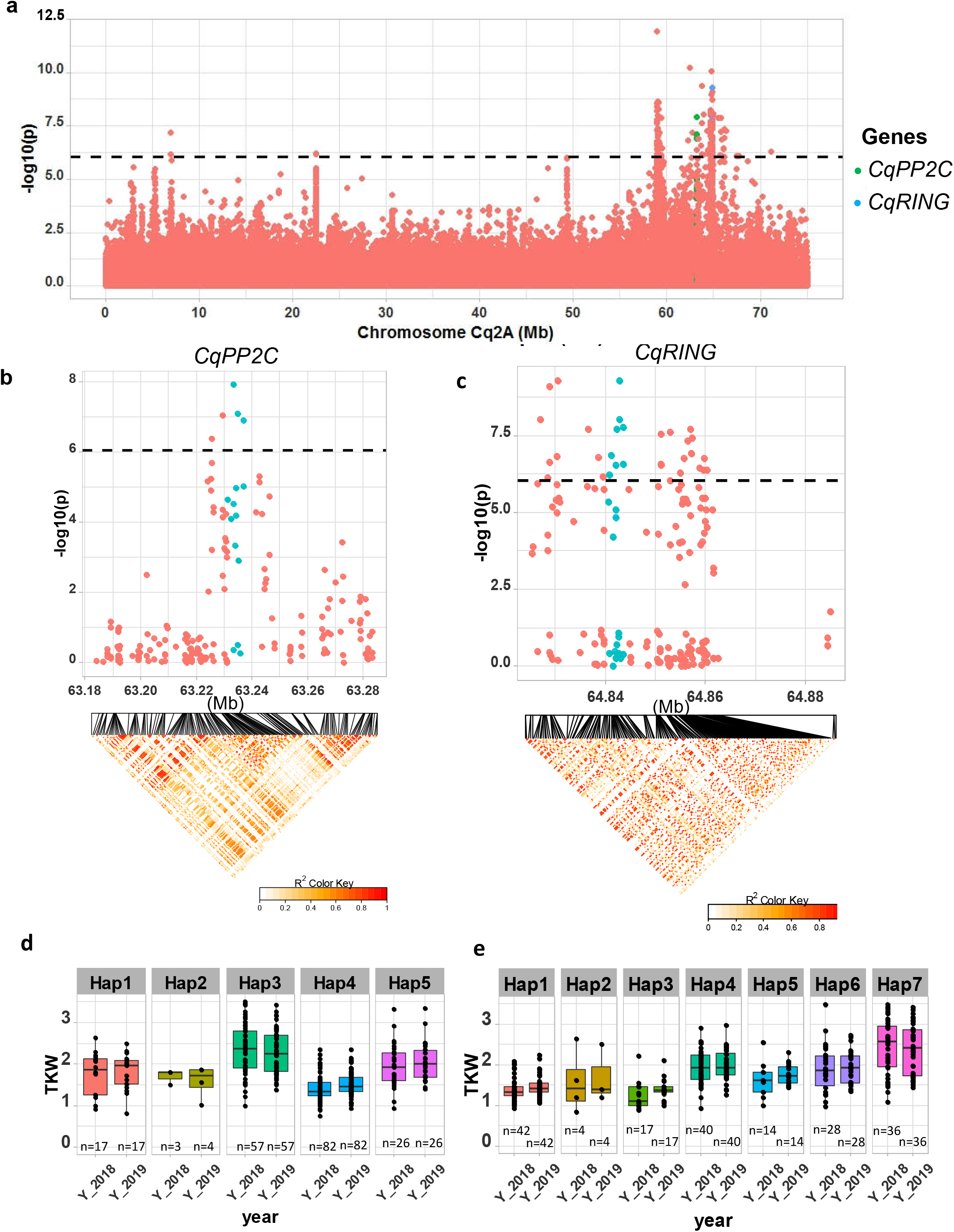
Identification of candidate genes for thousand seed weight. (a) Manhattan plot from chromosome Cq8B. Green and blue dots are depicting the *CqPP2C5* and the *CqRING* gene, respectively. (b) Top: Local Manhattan plot in the neighborhood of the *CqPP2C* gene. Bottom: LD heat map. (c) Top: Local Manhattan plot in the neighborhood of the *CqRING* gene. Bottom: LD heat map. Differences in thousand seed weight between five *CqPP2C* (d) and seven *CqRING* haplotypes (e).

Downy mildew is one of the major diseases in quinoa, which causes massive yield damage. Notably, our GWAS identified strong MTA for resistance against this disease. The most significant SNPs are located in subgenome A (Supplementary Fig. 12). Thus, the A-genome progenitor seems to be the donor of downy mildew resistance. We identified a candidate gene within a region 38.99 - 39.03 Mb on chromosome Cq2A, which showed the highest significant association (Supplementary Fig. 14C). This gene encodes a protein with an NBS-LRR (nucleotide-binding site leucine-rich repeat) domain often found in resistance gene analogs with a function against mildew infection ^36^.

## Discussion

We assembled a diversity set of 303 quinoa accessions and seven accessions from wild relatives. Plants were grown under northern European conditions, and agronomically important traits were studied. In total, 2.9 million SNPs were found after re-sequencing. We found substantial phenotypic and genetic variation. Our diversity set was structured into two highly diverged populations, and genomic regions associated for this diversity were localized. Due to a high marker density, candidate genes controlling qualitative and quantitative traits were identified. The high genetic diversity and rapid LD breakdown are reflecting the short breeding history of this crop.

We were aiming to assemble the first diversity set, which represents the genetic variation of this species. Therefore, we established a permanent resource that is genotypically and phenotypically characterized. We believe that this collection is important for future studies due to the following reasons: We observed substantial phenotypic variation for all traits and high homogeneity within accessions. Moreover, low or absent phenotypic variation within accessions demonstrates homogeneity as expected for a self-pollinating species. Therefore, the sequence of one plant is representative of the whole accession, which is important for the power of the GWAS.

Today, over sixteen thousand accessions of quinoa are stored *ex-situ* in seed banks in more than 30 countries ^37^. Despite the enormous diversity, only a few accessions have been genotyped with molecular markers. We found a clear differentiation into Highland and Lowland quinoa. In previous studies, five ecotypes had been distinguished: Valley type, Altiplano type, Salar type, Sea level type, and Subtropical type ^19^. Adaptation to different altitudes, tolerance to abiotic stresses such as drought and salt, and photoperiodic responses are the major factors determining ecotypes ^18^. In our study, we could further allocate the quinoa accessions to five Highland and two Lowland subpopulations. This demonstrates the power of high-density SNP mapping to identity finer divisions at higher *K*. The origin of accessions and ecotype differentiation could be meaningfully interpreted by combining the information from phylogenetic data and population structure. As we expected, North American accessions (accessions obtained from USDA) were clustering with Chilean accessions, suggesting sequence-based characterization of ecotypes would be more informative and reproducible. Moreover, high-density SNP genotyping unveiled the origin of unknown or falsely labeled gene bank accessions, as recently proposed by Milner, et al. ^38^. The geographical origin of 52 accessions from our panel was unknown. We suggest using phylogenic data and admixture results to complement the available passport data. For instance, two accessions with origin recorded as Chile are closely related to Peruvian and Bolivian accessions, which suggests that they are also originating from Highland quinoa.

What can we learn about the domestication of quinoa and its breeding history by comparing our results with data from other crops? LD decay is one parameter reflecting the intensity of breeding. LD decay in quinoa (32.4 kb) is faster than in most studies with major crop species, e.g. rapeseed (465.5 kb) ^39^, foxtail millet (*Setaria italica*, 100 kb) ^40^, pigeonpea (*Cajanus cajan*, 70 kb) ^41^, soybean (150 kb) ^42^ and rice (200 kb) ^43^. Although comparisons must be regarded with care due to different numbers of markers and accessions, different types of reproduction, and the selection intensity, the rapid LD decay in quinoa reflects its short breeding history and low selection intensity. Moreover, quinoa is a self-pollinating species where larger linkage blocks could be expected. However, cross-pollination rates in some accessions can be up to 17.36 % ^12^, which is exploited by small Andean farmers who grow mixed quinoa accessions to ensure harvest under different biotic and abiotic stresses. This may facilitate a certain degree of cross-pollination and admixture.

Interestingly, the LD structure between Highland and Lowland populations is highly contrasting (6.5 vs. 49.8 kb), indicating larger LD blocks in the Lowland population. Low nucleotide diversity and negative Tajima’s *D* were also observed in the Lowland population compared to Highland quinoa. The population differentiation index and LD differences have been used to test the hypothesis of multiple domestication events. As an example, different domestication bottlenecks have been reported for japonica (LD decay: 65 kb) and indica rice (LD decay: 200 kb) ^44^. The estimated *F*_*ST*_ value from this study (0.36) is in the similar range of *F*_*ST*_ estimates in rice subspecies *indica* and *japonica* (0.55) ^45^ and melon (*Cucumis melo*) subspecies *melo* and *agrestis* (0.46) ^46^. Two hypotheses have been proposed for the domestication of quinoa from *C. hircinum*; (1) one event that gave rise to Highland quinoa and subsequently to Lowland quinoa and (2) two separate domestication events giving rise to Highland and Lowland quinoa independently ^9^. However, our study is not strictly following the second hypothesis because *C. hircinum* accession BYU 566 was basal to both clades of the phylogenetic tree (Highland and Lowland). Moreover, our wild Chenopodium germplasm does not represent enough diversity for in-depth analysis of domestication events. Therefore, we propose three possible scenarios to explain strong differences in LD structure, nucleotide diversity, Tajima’s *D* and *F*_*ST*_, (1) two independent domestication evens with a strong bottleneck on lowland populations, (2) a single domestication but strong population growth after adaptation of lowland quinoa or (3) strong adaptive selection after domestication. To understand the history and genetics of domestication, it will be necessary to sequence a large representative set of outgroup species such as *C berlandieri*, *C. hircinum, C. pallidicaule,* and *C. suecicum.*

Apart from marker density and sample size, the power of GWAS depends on the quality of the phenotypic data. Plants were grown in Northern Europe. Therefore, the MTAs are, first of all, relevant for temperate long-day climates. The share of genetic variances and thus, the heritabilities were high across environments. We expect higher genotype x environment interaction for flowering time, days to maturity, plant height, and panicle length if short-day environments will be included because many accessions have a strong day-length response (data not shown). Furthermore, the positions of genes controlling Mendelian traits were precisely coinciding with significant SNP positions, as exemplified by the genes associated with saponin content and flower color. Hence, the diversity panel provides sufficient power to identify SNP-trait associations for important agronomic traits such as TSW and downy mildew tolerance. In different plant species, seed size is controlled by six different pathways ^35^. We found two important genes controlling seed size from the Brassinosteroid (*CqPP2C*) and the ubiquitin-proteasome (*CqRING*) pathway. The non-functional allele of soybean *PP2C1* resulted in small seeds ^34^. We detected a superior haplotype (PP2C_hap3), which results in larger seeds. *CqRING* encodes an E3 ubiquitin ligase protein. There are two RING-type E3 ubiquitins known as *DA1* and *DA2,* which are involved in seed size controlling pathway. They were found in Arabidopsis rice, maize, and wheat. Downy mildew is the most acute disease for quinoa, caused by the fungus *Peronospora variabili* ^47^. A recent study attempted identification of genes based on a GWAS analysis. However, no significant associations were found, probably due to the lack of power because of the small number of accessions used (61 and 88) ^48^. In our study, a strong MTA suggests that the NBS-LRR gene on chromosome Cq2A contributes to downy mildew resistance in quinoa. We propose using this sequence for marker-assisted selection in segregating F2 populations produced during pedigree breeding of quinoa.

In this study, the advantage of multivariate analysis of cross-phenotype association became obvious. We could identify candidate genes with a pleiotropic effect on days to flowering, days to maturity, plant height, and panicle length. Interestingly, the most significant SNP was residing within a putative *GLX-2* ortholog. *GLX* genes, among other functions, have been shown to impact cell division and proliferation in *Amaranthus paniculatus* ^49^. Therefore, the *CqGLX-2* gene is one candidate for controlling day length response.

This study also has a major breeding perspective. We aimed to elucidate the potential of quinoa for cultivation in temperate climates. Evidently, many accessions are not adapted to northern European climate and photoperiod conditions because they flowered too late and did not reach maturity before October. Nevertheless, 48 accessions are attractive as crossing partners for breeding programs because they are insensitive to photoperiod or long-day responsive. Moreover, they are attractive due to their short plant height, low tillering capacity, favorable inflorescence architecture, and high TSW. These are important characters for mechanical crop cultivation and combine harvesting. The MTA found in this study offers a perspective to use parents with superior phenotypes in crossing programs. We suggest a genotype building strategy by pyramiding favorable alleles (haplotypes). In this way, also accessions from our diversity set, which are not adapted to long-day conditions but with favorable agronomic characters, will be considered. Then, favorable genotypes will be identified from offspring generations by marker-assisted selection using markers in LD with significant SNPs. Furthermore, the MTA from this study will be useful for allele mining in quinoa germplasm collections to identify yet unexploited genetic variation.

## Materials and Methods

### Plant materials and growth conditions

We selected 350 quinoa accessions for phenotyping, and of these, 296 were re-sequenced in this study. Re-sequencing data of 14 additional accessions that had already been published ^9^ were also included in the study, together with the wild relatives (*C. belandieri* and *C. hircinum*) ^9^. These accessions represent different geographical regions of quinoa cultivation (Supplementary Table 1). Plants were grown in the field in Kiel, Northern Germany, in 2018 and 2019. Seeds were sown in the second week of April in 35x multi-tray pots. Then plants were transplanted to the field in the first week of May as single-row plots in a randomized complete block design with three blocks. The distances between rows and between plants were set to 60 cm and 20 cm, respectively. Each row plot contained seven plants per accession.

We recorded days to bolting (DTB) as BBCH51 and days to flowering (DTF) as BBCH60 twice a week during the growth period. Days to maturity (DTM) was determined when plants reached complete senescence (BBSHC94). If plants did not reach this stage, DTM was set as 250 days. In both years, plants were harvested in the second week of October. Plant height (PH), panicle length (PL), and the number of branches (NoB) were phenotyped at harvest. Stem lying (STL) (Supplementary Fig. 2) was scored on a scale from one to five, where score one indicates no stem lying. Similarly, panicle density was recorded on a scale from one to seven, where density one represents lax panicles, and panicle density seven represents highly dense panicles. Flower color and stem color were determined by visual observation. Pigmented and non-pigmented plants were scored as 1 and 0, respectively. Growth type was classified into two categories and analyzed as a dichotomous trait as well. We observed severe mildew infection in 2019. Therefore, we scored mildew infection on a scale from 1 to 3, where 1 equals no infection, and 3 equals severe infection.

### Statistical analysis

We calculated the best linear unbiased estimates of the traits across years by fitting a linear mixed model using the lme4 R package ^50^. We used the following model:

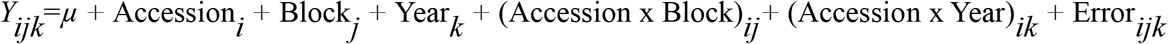

Where *μ* is the mean, Accession*i* is the genotype effect of the *i*-th accession, Block*j* is the effect of the *j*-th Block, Year*k* is the effect of the *k*-th year, (Accession x Block)_*ij*_ is the Accession-Block interaction effect, Accession x Year_*ik*_ is the accession-year interaction effect, Error_*ijk*_ is the error of the *j*-th block in the *k*-th year. We treated all items as random effects for heritability estimation, and for best linear unbiased estimates (BLUE), accessions were treated as fixed effects. We analyzed the principle components of phenotypes using the R package FactoMineR ^51^.

### Genome sequencing and identification of genomic variations

For DNA extraction, two plants per genotype were grown in a greenhouse at the University of Hohenheim, and two leaves from a single two-months old plant were collected and frozen immediately. DNA was subsequently extracted using the AX Gravity DNA extraction kit (A\&A Biotechnology, Gdynia, Poland) following the manufacturer’s instructions. Purity and quality of DNA were controlled by agarose gel electrophoresis and the concentration determined with a Qubit instrument using SYBR green staining. Whole-genome sequencing was performed for 312 accessions at Novogene (China) using short-reads Illumina NovaSeq S4 Flowcell technology and yielded an average of 10 Gb of paired-end (PE) 2 × 150 bp reads with quality Q>30 Phred score per sample, which is equivalent to ~7X coverage of the haploid quinoa genome (~1.45 Gb). We then used an automated pipeline (https://github.com/IBEXCluster/IBEX-SNPcaller/blob/master/workflow.sh) compiled based on the Genome Analysis Toolkit. Raw sequence reads were filtered with trimmomatic-v0.38 ^52^ using the following criteria: LEADING:20; TRAILING:20; SLIDINGWINDOW:5:20; MINLEN:50. The filtered paired-end reads were then individually mapped for each sample against an improved version of the QQ74 quinoa reference genome (CoGe id53523) using BWA-MEM (v-0.7.17) ^53^ followed by sorting and indexing using samtools (v1.8) ^54^. Duplicated reads were marked, and read groups were assigned using the Picard tools (http://broadinstitute.github.io/picard/). Variants were identified with GATK (v4.0.1.1) ^55 56^ using the “--emitRefConfidence” function of the HaplotypeCaller algorithm and “—heterozygosity” value set at 0.005 to call SNPs and InDels for each accession. Individual g.vcf files for each sample were then compressed and indexed with tabix (v-0.2.6) ^57^ and combined into chromosome g.vcf using GenomicsDBImport function of GATK. Joint genotyping was then performed for each chromosome using the function GenotypeGVCFs of GATK. To obtain high confidence variants, we excluded SNPs with the VariantFiltration function of GATK with the criteria: QD < 2.0; FS > 60.0; MQ < 40.0; MQRankSum < −12.5; ReadPosRankSum < −8.0 and SOR > 3.0. Then, SNP loci which contained more than 70% missing data, were filtered by VCFtools ^58^ (v0.1.5), which resulted in our initial set of ~45M SNPs for all the 332 accessions, including 20 previously re-sequenced accessions ^9^. All resequencing data are submitted to SRA under project id BioProject PRJNA673789.

In our panel, we had three triplicates for quality checking and nine duplicates between Jarvis et al. 2017 and 312 newly re-sequenced accessions. In order to remove duplicates, as a preliminary analysis, we removed SNP loci with a minimum mean-depth <5 across samples and SNP loci with more than 5% missing data. Then, we filtered SNPs with a minor allele frequency lower than 0.05 (MAF<0.05). After these filtering steps, we obtained a VCF file that contained 229,017 SNPs. Then, we construct a maximum likelihood (ML) tree. First, we used the modelFinder ^59^ in IQ-TREE v1.6.619 (Nguyen et al. 2015) to determine the best model for ML tree construction. We selected GTR+F+R8 (GTR: General time-reversible, F: Empirical base frequencies, R8: FreeRate model) as the best fitting model according to the Bayesian Information Criterion (BIC) estimated by the software. We used 1000 replicates with ultrafast bootstrapping (UFboots)^60^ to check the reliability of the phylogenetic tree. To visualize the phylogenetic tree, we used the Interactive Tree Of Life tool (https://itol.embl.de/)^61^. Then, based on the phylogenetic tree, we removed duplicate accessions and accessions with unclear identity. After the quality control, we retained 310 accessions (303 quinoa accessions and 7 wild *Chenopodium* accessions).

Then we used the initial SNP set and defined two subsets using the following criteria: (1) A base SNP set of 5,817,159 biallelic SNPs obtained by removing SNPs with more than 50% missing genotype data, minimum mean depth less than five, and minor allele frequency less than 1%. (2) A high confidence (HCSNP) set of 2,872,935 SNPs from the base SNP set by removing SNPs with a minor allele frequency of less than 5%. The base SNP set was used for the diversity statistics, and the HCSNPs set was used for GWAS analysis.

We annotated the HCSNP using SnpEff 4.3T ^27^ and a custom database ^27^ based on the QQ74 reference genome and annotation (CoGe id53523). Afterward, we extracted the SNP annotations using SnpSift ^62^. Based on the annotations, SNPs were mainly categorized into five groups, (1) upstream of the transcript start site (5kb), (2) downstream of the transcript stop site (5kb), (3) coding sequence (CDS), (4) intergenic, and (5) intronic. We used SnpEff to categorize SNPs in coding regions based on their effects such as synonymous, missense, splice acceptor, splice donor, splice region, start lost, start gained, stop lost, and spot retained.

### Phylogenetic analysis and population structure analysis

For population structure analysis, we employed SNP subsets, as demonstrated in previous studies, to reduce the computational time ^63^. We created ten randomized SNP sets, each containing 50,000 SNPs. To create subsets, first, the base SNP set was split into 5000 subsets of an equal number of SNPs. Then, 10 SNPs from each subset were randomly selected, providing a total of 50,000 SNPs in a randomized set (randomized 50k set). We then repeated this procedure for nine more times and finally obtained ten randomized 50k sets. Population structure analysis was conducted using ADMIXTURE (Version: 1.3) ^64^. We ran ADMIXTURE for each subset separately with a predefined number of genetic clusters K from 2 to 10 and varying random seeds with 1000 bootstraps. Also, we performed the cross-validation (CV) procedure for each run. Obtained Q matrices were aligned using the greedy algorithm in the CLUMPP software ^65^. Population structure plots were created using custom R scripts. We then combined SNP from the ten subsets to create a single SNP set of 434,077 unique SNPs for the phylogenetic analysis. We used the same method mentioned above to create the phylogenetic tree. Here we selected the model GTR+F+R6 based on the BIC estimates. For the principal component analysis (PCA) we used the HCSNP set and analysis was done in R package SNPrelate ^66^. We estimated the top 10 principal components. The first (PC1) and second (PC2) were plotted using custom R scripts.

### Genomic patterns of variations

Using the base SNP set, we calculated nucleotide diversity (π) for subpopulations and π ratios for Highland and Lowland population regions with the top 1% ratios of π Highland/ π Lowland candidate regions for population divergence. We also estimated Tajima’s *D* values for both populations to check the influence of selection on populations. *F*_*ST*_ values were calculated between Highland and Lowland populations using the 10kb non-overlapping window approach. Nucleotide diversity, Tajima’s *D*, and *F*_*ST*_ calculations were carried out in VCFtools (v0.1.5) ^58^.

### Linkage disequilibrium analysis

First, we calculated linkage disequilibrium in each population separately (Highland and Lowland). Then, LD was calculated in the whole population, excluding wild accessions. For LD calculations, we further filtered the HCSNP set by removing SNPs with >80% missing data ^29^. Using a set of 2,513,717 SNPs, we calculated the correlation coefficient (*r*^*2*^) between SNPs up to 300kb apart by setting -MaxDist 300 and default parameters in the PopLDdecay software ^67^. LD decay was plotted using custom R scripts based on the ggplot2 package.

### Genome-wide association study

We used the best linear unbiased estimates (BLUE) of traits and HCSNPs for the GWAS analysis. Morphological traits were treated as dichotomous traits and analyzed using generalized mixed linear models with the lme4 R software package ^50^. We used population structure and genetic relationships among accessions to minimize false-positive associations. Population structure represented by the PC was estimated with the SNPrelate software ^66^. Genetic relationships between accessions were represented by a kinship matrix calculated with the efficient mixed-model association expedited (EMMAX) software ^68^ using HCSNPs. Then, we performed an association analysis using the mixed linear model, including K and P matrices in EMMAX. We estimated the effective number of SNPs (n=1,062,716) using the Genetic type I Error Calculator (GEC) ^69^. We set the significant *P*-value threshold (Bonferroni correction, 0.05/n, −log10(4.7e-08)=7.32) and suggestive significant threshold (1/n, −log10(9.41e-7)= 6.02) to identify significant loci underlying traits. We plotted SNP *P*-values on Manhattan plots using the qqman R package ^70^.

## Acknowledgments

We thank David Jarvis for providing the updated version of the quinoa reference genome. We thank Monika Bruisch, Brigitte Neidhardt-Olf, Elisabeth Kokai-Kota, Verena Kowalewski, and Gabriele Fiene for technical assistance. The financial support of this work was provided by the Competitive Research Grant (Grant No. OSR-2016-CRG5-466 2966-02) of the King Abdullah University of Science and Technology, Saudi Arabia and baseline funding from KAUST to Mark Tester.

## Author contributions

C.J, M.T, and N.E directed the project and conceived the research. D.S.R.P conducted genomic data analysis and GWAS analysis. D.S.R.P and N.E performed field experiments and phenotyping. E.R conducted SNP identifications. G.W and S.M.S selected and assembled the diversity panel. K.S contributed to DNA isolation, library preparation for genome sequencing. D.S.R.P, together with all authors, wrote and finalized the manuscript.

## Data availability

The raw sequencing data have been submitted to the NCBI Sequence Read Archive (SRA) under the BioProject PRJNA673789. Quinoa reference genome version 2 is available at CoGe database under genome id 53523. Source data are provided with the paper.

## Code availability

Custom scripts used for SNP calling are available on GitHub: https://github.com/IBEXCluster/IBEX-SNPcaller/blob/master/workflow.sh. Additional information on other custom scripts will be available upon request.

## Supplementary data

### Supplementary tables

**Supplementary Table 1:** Accessions from the quinoa diversity panel and results from re-sequencing

**Supplementary Table 2:** Summary of high-quality SNPs identified in quinoa accessions

**Supplementary Table 3:** Variance components analysis of 12 quantitative traits

**Supplementary Table 4**: Summary of marker trait associations (MTA)

**Supplementary Table 5:** Candidate genes linked to SNP with significant trait associations

**Supplementary Table 6:** Summary of MTA associated with DTF, DTM, PD and PH identified on chromosome Cq2A

**Supplementary Table 7:** Candidate genes located within the 50kb flanking regions of significantly associated SNPs from the multivariate GWAS analysis

### Supplementary figures

**Supplementary Fig. 1:** Geographical origin of the accessions forming the quinoa diversity panel.

**Supplementary Fig. 2:** Overview of the field experiment and exemplary images demonstrating phenotypic traits; (A) and (B): Overview of the field and phenotypic variation among accession; (C): Bolting (BBCH51) and (D) flowering (BBCH60) stage; Glomerulate (E) and amarantiform (F) panicle shapes; red (G) and green (H) stem color; red (I) and green (J) flower/inflorescence; Growth type 1 (K) and type 5 (L); (M): Plant height and maturity variation between two accessions.

**Supplementary Fig. 3:** SNP density heat map across the 18 quinoa chromosomes. Different colors depict SNP density.

**Supplementary Fig. 4:** Chromosome wide LD decay in genome A (A) and genome B (B). Colors are depicting different chromosomes. (C) Genome-wide average LD decay of the A sub-genome (blue) and B sub-genome (red).

**Supplementary Fig. 5:** SNP based PCA across all 18 quinoa chromosomes. Red circles are depicting the two clusters of Lowland accessions.

**Supplementary Fig. 6** (A) ADMIXTURE ancestry coefficients for K ranging from 3 to 7 and 9. Each vertical bar represents an accession, and color proportions on the bar correspond to the genetic ancestry. (B) Cross-validation error in ADMIXTURE run.

**Supplementary Fig. 7**: Diversity of populations along chromosomes measured based on 10 kb non-overlapping windows. Nucleotide diversity (π) distribution of 10 kb windows in population Highland (A) and Lowland (B). (C) Nucleotide diversity ratios (π Lowland/ π Highland). (D) Pairwise genome-wide fixation index (*F*_*ST*_) between Highland and Lowland. The broken horizontal line represents the top 1% threshold.

**Supplementary Fig. 8:** Distribution of Tajima’s *D* along chromosomes in Highland (B) and Lowland (D) populations. Density distribution of Tajima’s *D* between populations. Different colors represent the quartiles.

**Supplementary Fig. 9:** Graphical presentation of correlations between years among 12 traits. Pearson correlation value (*R*) with *P*-values are shown. DTB: days to bolting (inflorescence emergence), DTF: days to flowering, DTB to DTF: days between bolting and flowering, DTM; days to maturity, PH: plant height (cm), PL: panicle length (cm), PD: panicle density (cm), NoB: Number of branches, STL: stem lying, Saponin: saponin content as foam height (mm), Seed yield: seed yield per plant (g), TSW: thousand seed weight (g),

**Supplementary Fig. 10:** Pearson correlations among 12 quinoa traits. Best linear unbiased estimates across two years were used. Below the diagonal, scatter plots are shown with the fitted line in red. Above the diagonal, the Pearson correlation coefficients are shown with significance levels, *** =*P*<0.001, **=*P*<0.01.

**Supplementary Fig. 11:** PCA of 12 quantitative phenotypes. A: Individual factor map colored according to populations identified from SNP analysis. B: Variables factor map of the PCA.

**Supplementary Fig. 12:** Manhattan plots from GWAS with data from 2018 (left), 2019 (center), and the mean of both years (right): The blue horizontal line indicates the suggestive threshold - log_10_ (8.98E-7). The red horizontal line indicates the significant threshold (Bonferroni correction) - log_10_(1.67e-8).

**Supplementary Fig. 13:** Quantile-quantile plots of GWAS in two years, 2018 (left) and 2019 (center), and BLUE (right).

**Supplementary Fig. 14:** Local Manhattan plots for (A) flower color, (B) saponin content, and (C) mildew infection. Candidate genes are shown in the color legend. LD heat maps are placed at the Bottom. The colors of the heat map represent the pairwise correlation between individual SNPs.

**Supplementary Fig. 15:** PCA of 4 quantitative traits (DTF, DTM, PH, and PL). A: Individual factor map, B: variables factor map of the PCA, C: distribution of the first three principal components which were used for GWAS analysis.

**Supplementary Fig. 16:** GWAS analysis of principal components, PC1 (A), PC2 (B), PC3 (C): Manhattan plots (left), and quantile-quantile plots (right): The blue horizontal line in the Manhattan plots indicates the suggestive threshold −log_10_(8.98E-7). The red horizontal line indicates the significance threshold (Bonferroni correction) −log_10_(1.67e-8).

**Supplementary Fig. 17:** Haplotypes of two genes, *CqPP2C* and *CqRING* controlling seed size in quinoa. Geographic origin of the accessions and haplotype networks are displayed below the gene structure.

