## Supplementary Fig for "Genome-wide association study in the pseudocereal quinoa reveals selection pattern typical for crops with a short breeding history"

### Supplementary tables

**Supplementary Table 1**: Summary of marker trait associations (MTA)

| Trait | No. of significant MTA 2018 | No. of significant MTA 2019 | No. of significant MTA BLUE | No. of consistant MTA |
|---|---|---|---|---|
| DTB | 23 | 3 | 9 | 2 |
| DTB to DTF | 4 | 3 | 5 | 0 |
| DTF | 18 | 26 | 25 | 13 |
| DTM | 640 | 680 | 674 | 387 |
| PH | 43 | 23 | 54 | 11 |
| PL | 13 | 13 | 26 | 2 |
| NoB | 49 | 0 | 8 | 0 |
| PD | 91 | 11 | 46 | 2 |
| TKW | 316 | 496 | 508 | 168 |
| Yield | 7 | 49 | 19 | 0 |
| Saponin | 17 | 36 | 44 | 6 |
| STL | 13 | 1 | 4 | 0 |
| FC | 10 | 4 | 10 | 4 |
| GT | 1 | 27 | 14 | 0 |
| SC | 3 | 6 | 6 | 3 |
| PSH | 27 | 90 | 28 | 2 |
| Mildew | NA | 86 | NA | NA |
| Total | 1275 | 1554 | 1480 | 600 |

**Supplementary Table 2:** Summary of MTA associated with DTF, DTM, PD and PH identified on chromosome Cq2A

| Chromosome | Position | Marker | DTF | DTM | PH | PL |
|---|---|---|---|---|---|---|
| | | | ***P-value*** | | | |
| Cq2A | 4242017 | 02:4242017 | 3.15E-08 | 6.85E-10 | 2.91E-08 | 4.43E-07 |
| Cq2A | 4406829 | 02:4406829 | 1.63E-08 | 8.56E-10 | 5.91E-08 | 2.83E-07 |
| Cq2A | 8080505 | 02:8080505 | 4.52E-07 | 2.01E-11 | 3.69E-08 | 8.20E-08 |
| Cq2A | 8080510 | 02:8080510 | 5.13E-07 | 1.83E-11 | 7.21E-08 | 7.58E-08 |
| Cq2A | 8093547 | 02:8093547 | 4.66E-09 | 3.96E-14 | 1.42E-08 | 1.50E-07 |
| Cq2A | 8101350 | 02:8101350 | 4.08E-07 | 4.06E-12 | 9.25E-08 | 1.46E-07 |
| Cq2A | 8116642 | 02:8116642 | 3.87E-08 | 3.43E-09 | 1.88E-08 | 7.67E-07 |

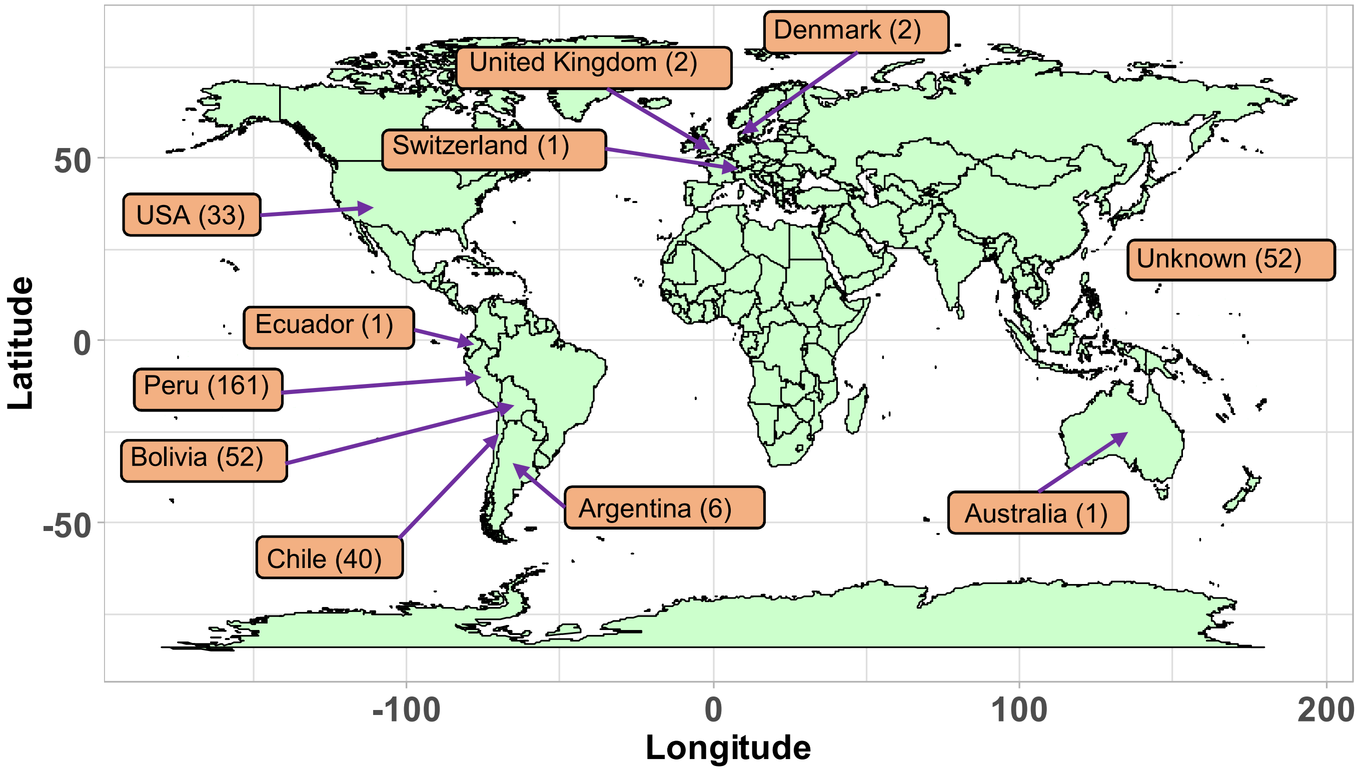


**Supplementary Fig. 1:** Geographical origin of the accessions forming the quinoa diversity panel.

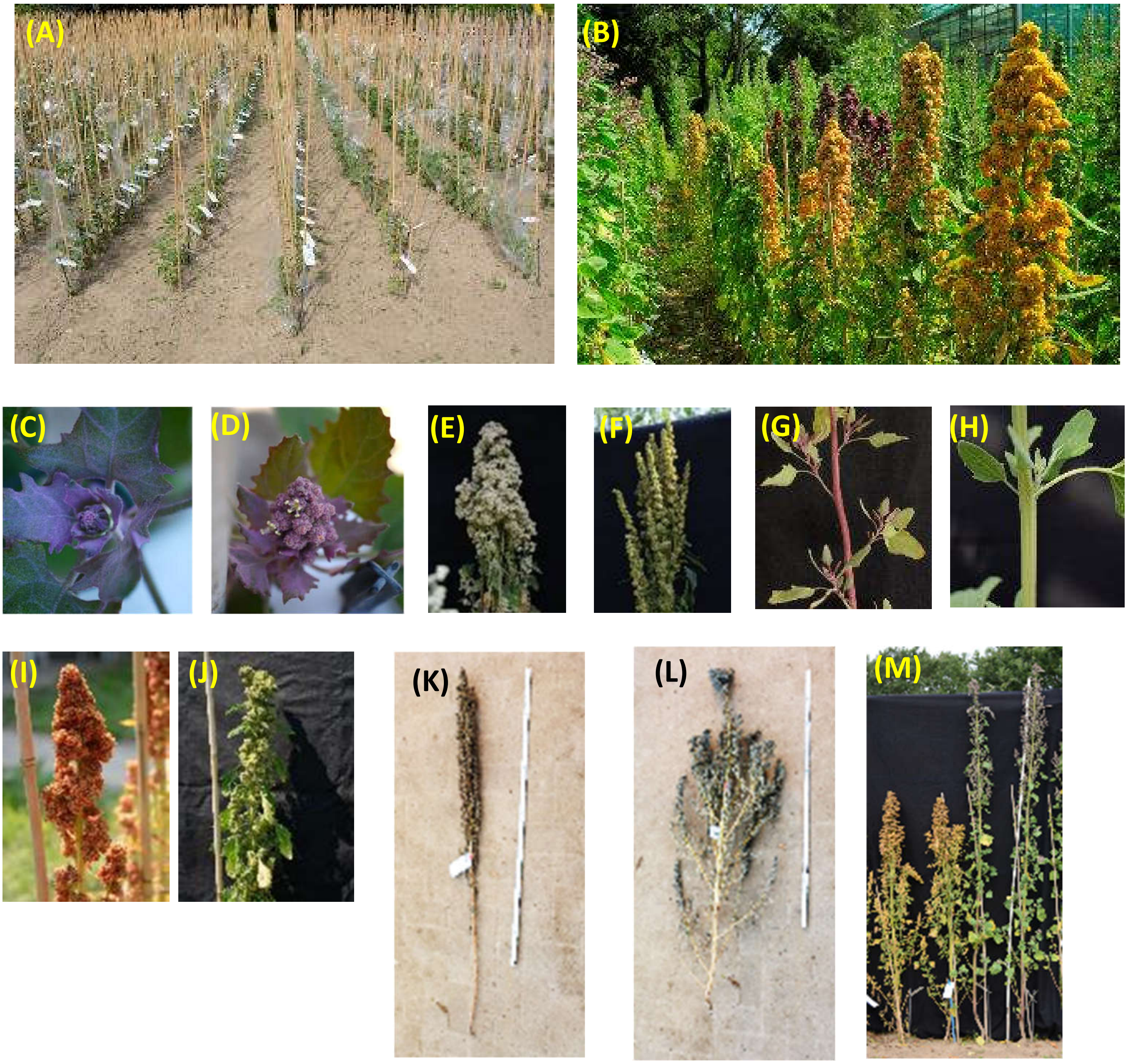


**Supplementary Fig. 2:** Overview of the field experiment and exemplary images demonstrating phenotypic traits; (A) and (B): Overview of the field and phenotypic variation among accession; (C): Bolting (BBCH51) and (D) flowering (BBCH60) stage; Glomerulate (E) and amarantiform (F) panicle shapes; red (G) and green (H) stem color ; red (I) and green (J) flower/inflorescence; Growth type 1 (K) and type 5 (L); (M): Plant height and maturity variation between two accessions.

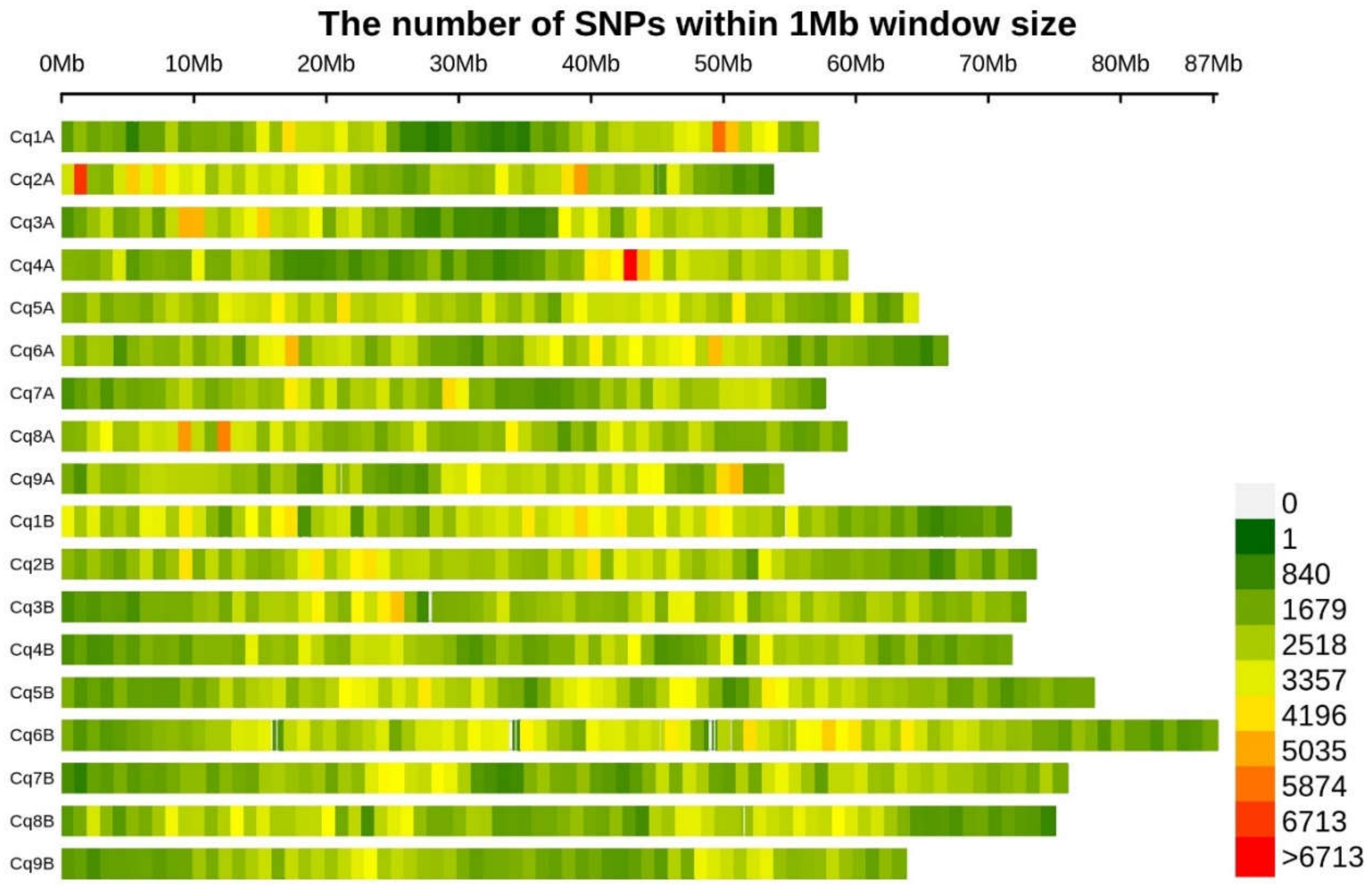


**Supplementary Fig. 3:** SNP density heat map across the 18 quinoa chromosomes. Different colors depict SNP density.

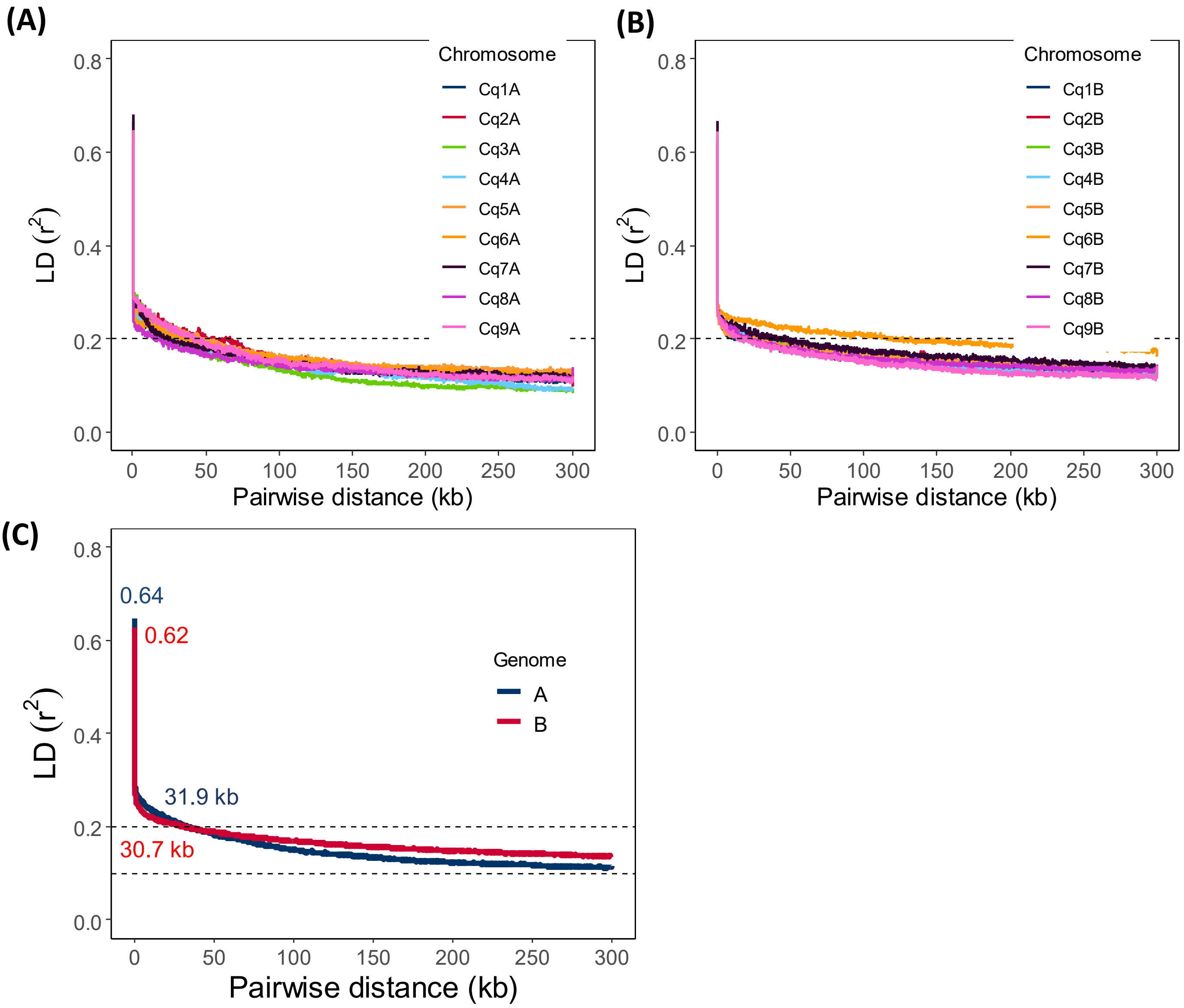


**Supplementary Fig. 4:** Chromosome wide LD decay in genome A (A) and genome B (B). Colors are depicting different chromosomes. (C) Genome-wide average LD decay of the A sub-genome (blue) and B sub-genome (red).

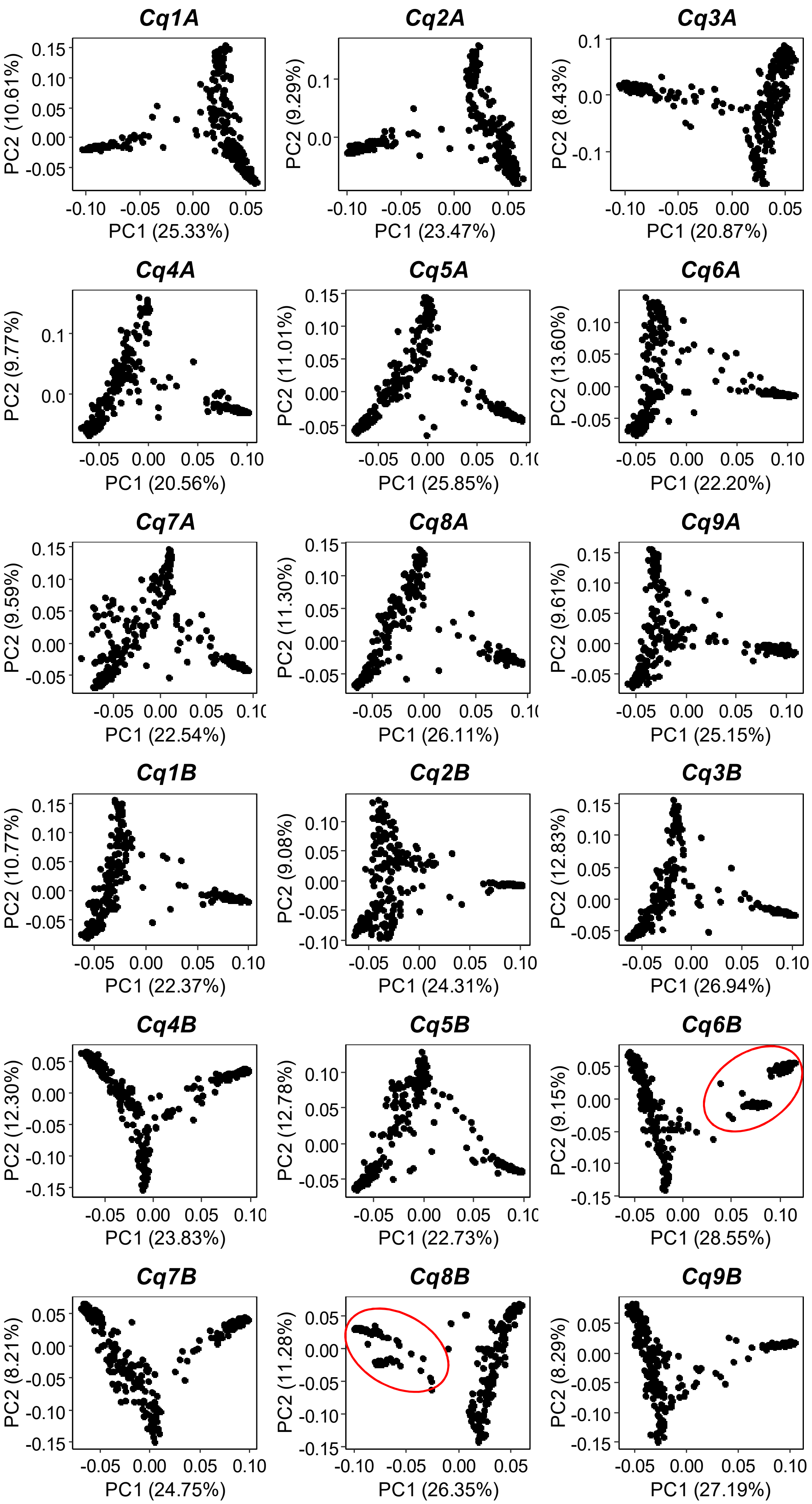


**Supplementary Fig. 5:** SNP based PCA across all 18 quinoa chromosomes. Red circles are depicting the two clusters of Lowland accessions.

**(A)**

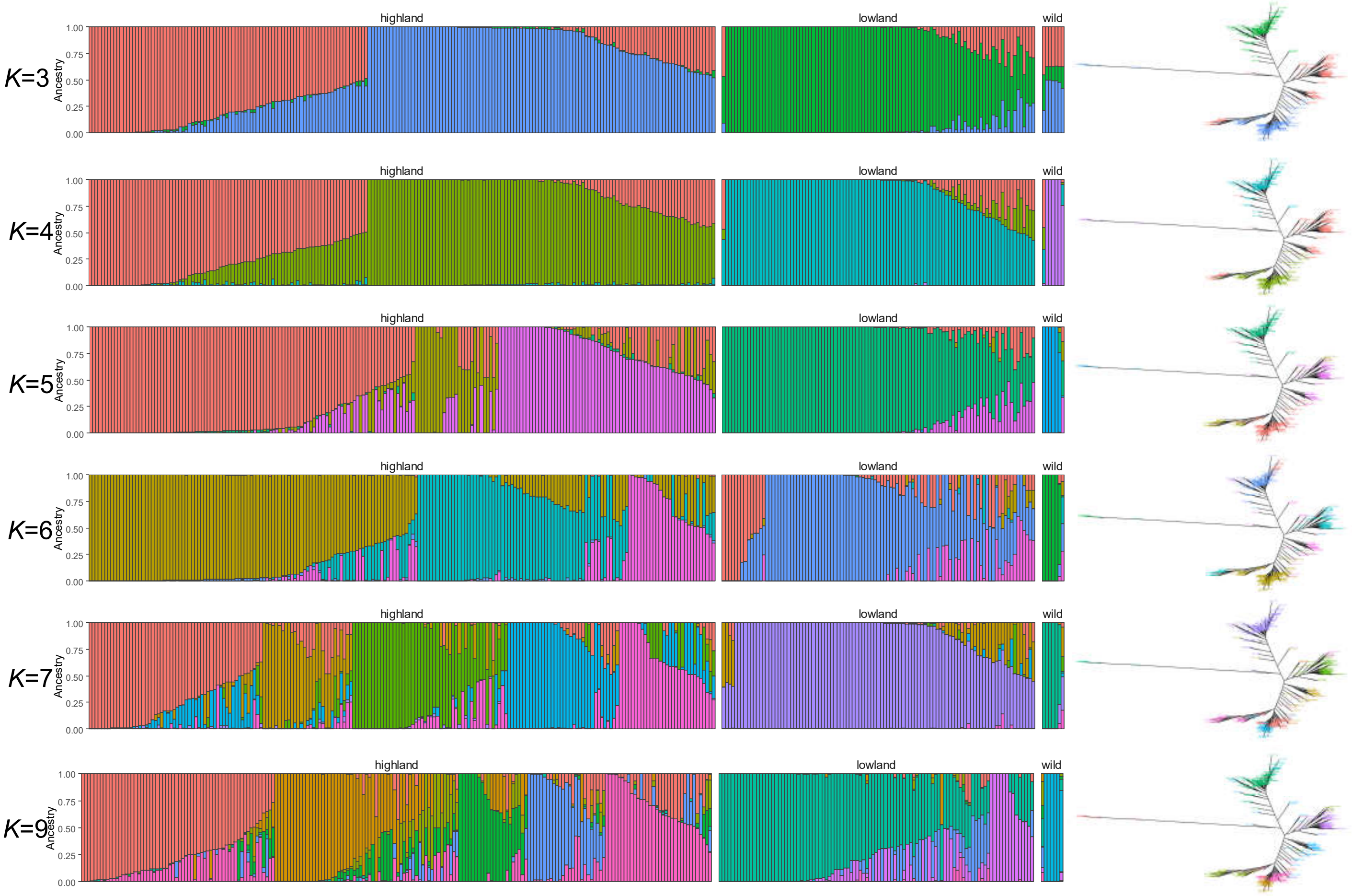


**(B)**

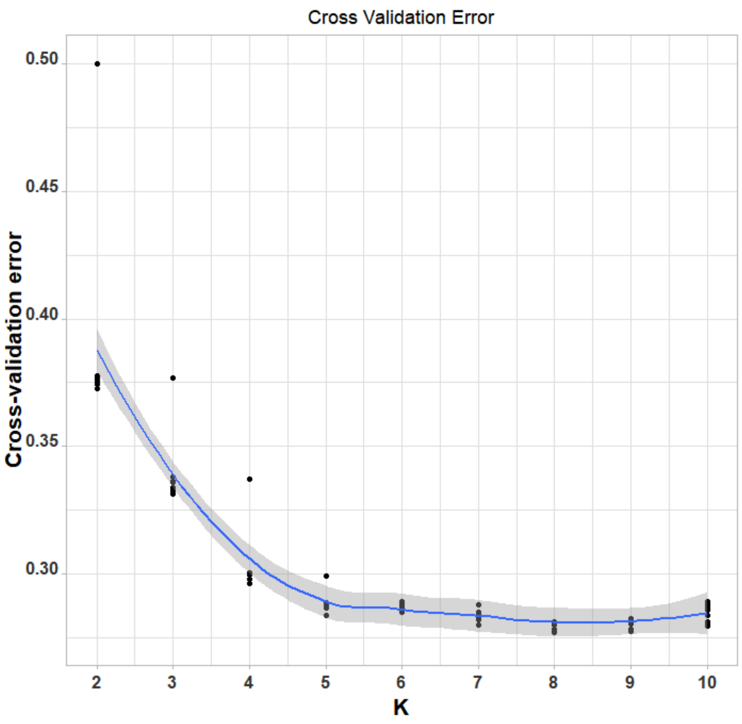


**Supplementary Fig. 6:** (A) ADMIXTURE ancestry coefficients for K ranging from 3 to 7 and 9. Each vertical bar represents an accession, and color proportions on the bar correspond to the genetic ancestry. (B) Cross-validation error in ADMIXTURE run.

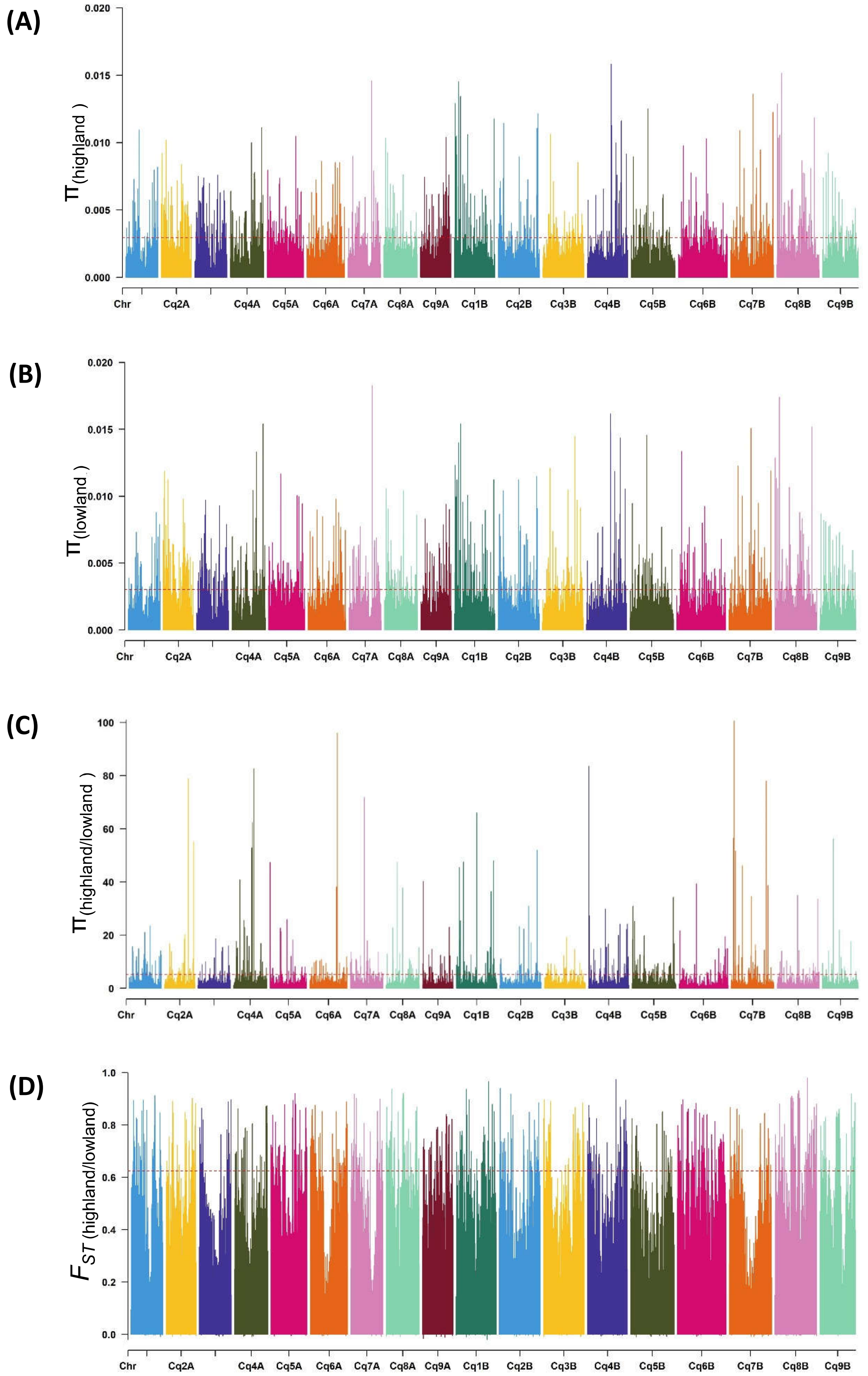


**Supplementary Fig. 7:** Diversity of populations along chromosomes measured based on 10 kb non-overlapping windows. Nucleotide diversity (π) distribution of 10 kb windows in population Highland (A) and Lowland (B). (C) Nucleotide diversity ratios (π Lowland/ π Highland). (D) Pairwise genome-wide fixation index ($F_{ST}$) between Highland and Lowland. The broken horizontal line represents the top 1% threshold.

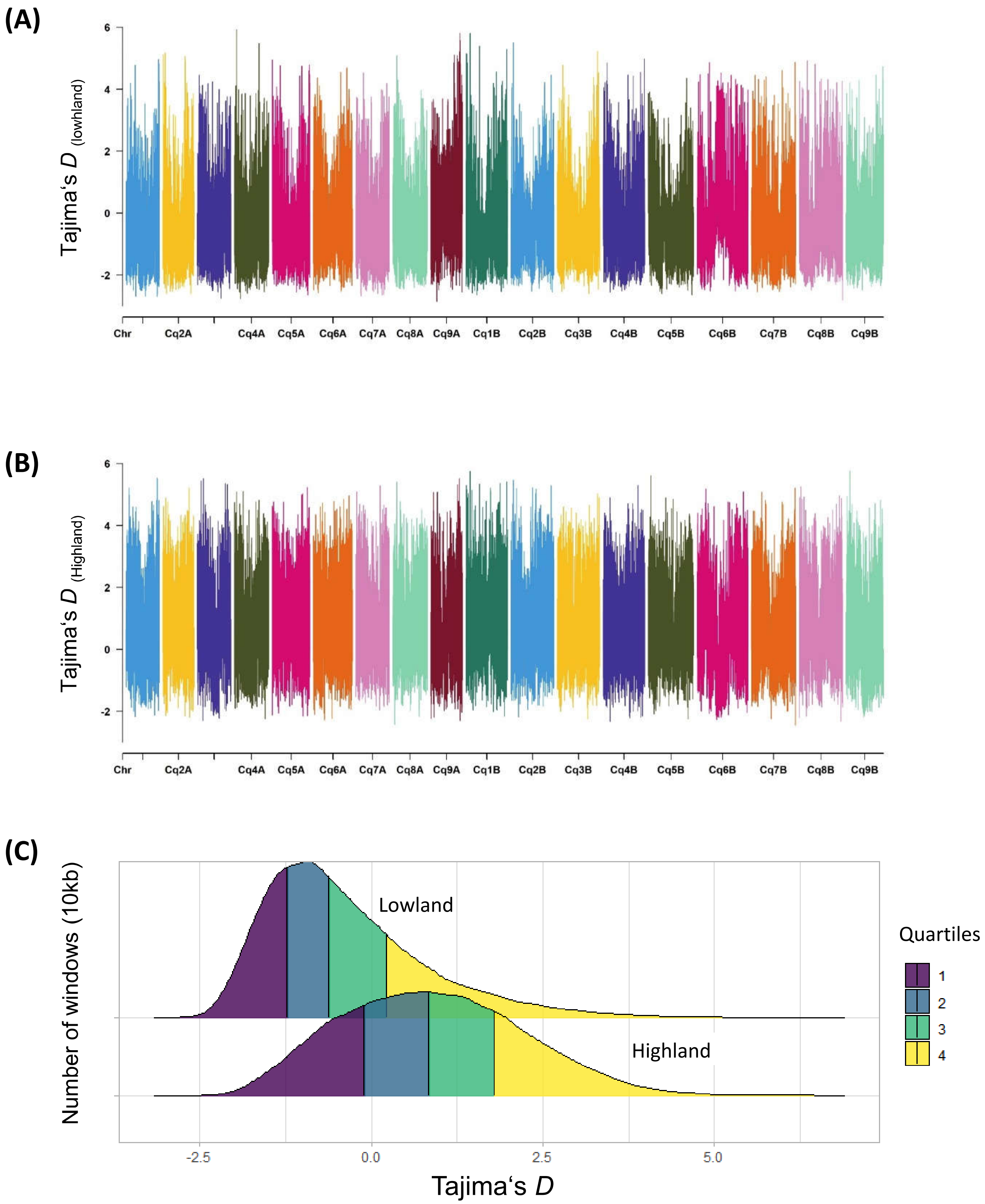


**Supplementary Fig. 8:** Distribution of Tajima's *D* along chromosomes in Highland (B) and Lowland (D) populations. Density distribution of Tajima's *D* between populations. Different colors represent the quartiles.

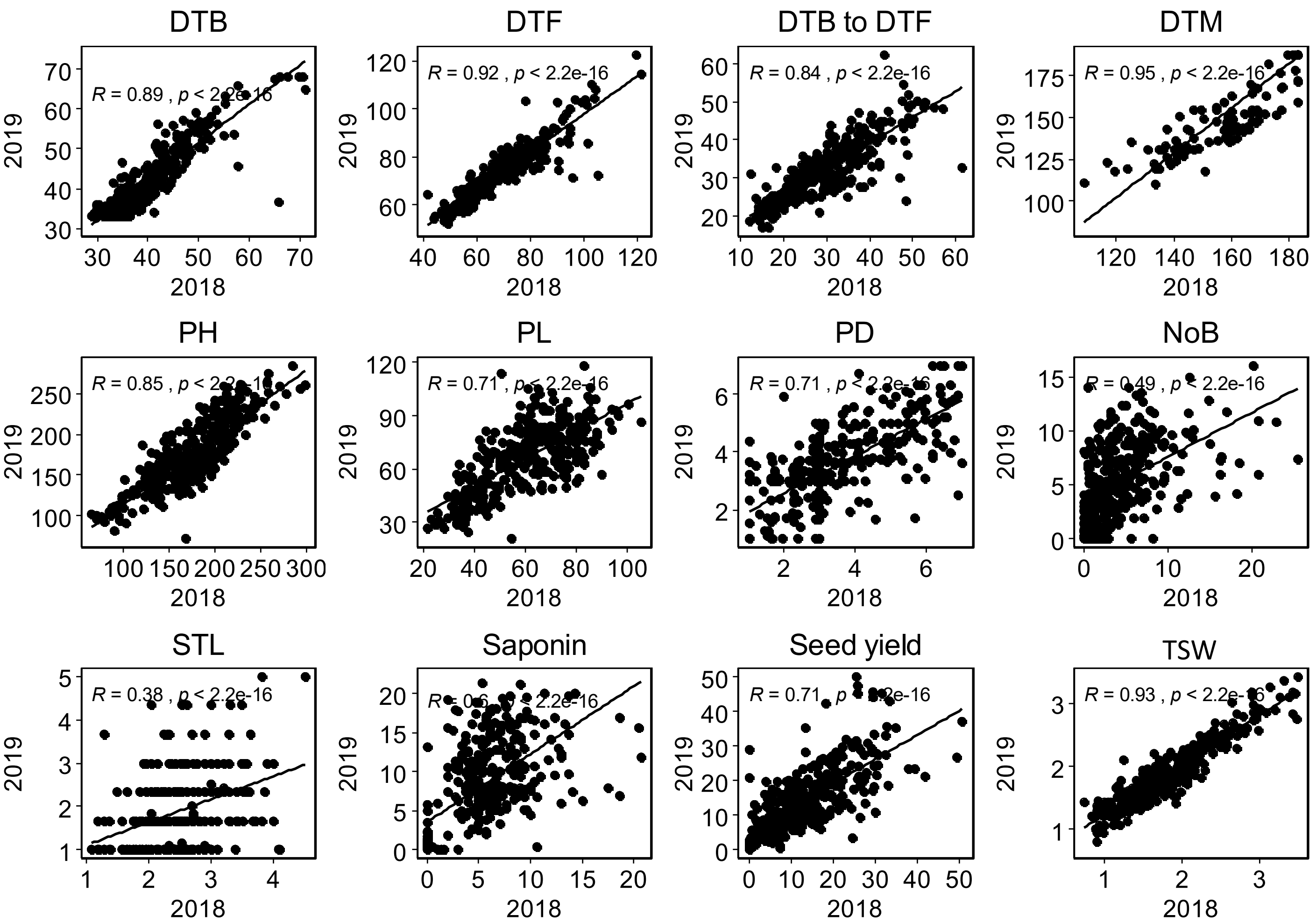


**Supplementary Fig. 9:** Graphical presentation of correlations between years among 12 traits. Pearson correlation value (R) with P-values are shown. DTB: days to bolting (inflorescence emergence), DTF: days to flowering, DTB to DTF: days between bolting and flowering, DTM; days to maturity, PH: plant height (cm), PL: panicle length (cm), PD: panicle density (cm), NoB: Number of branches, STL: stem lying, Saponin: saponin content as foam height (mm), Seed yield: seed yield per plant (g), TSW: thousand seed weight (g)

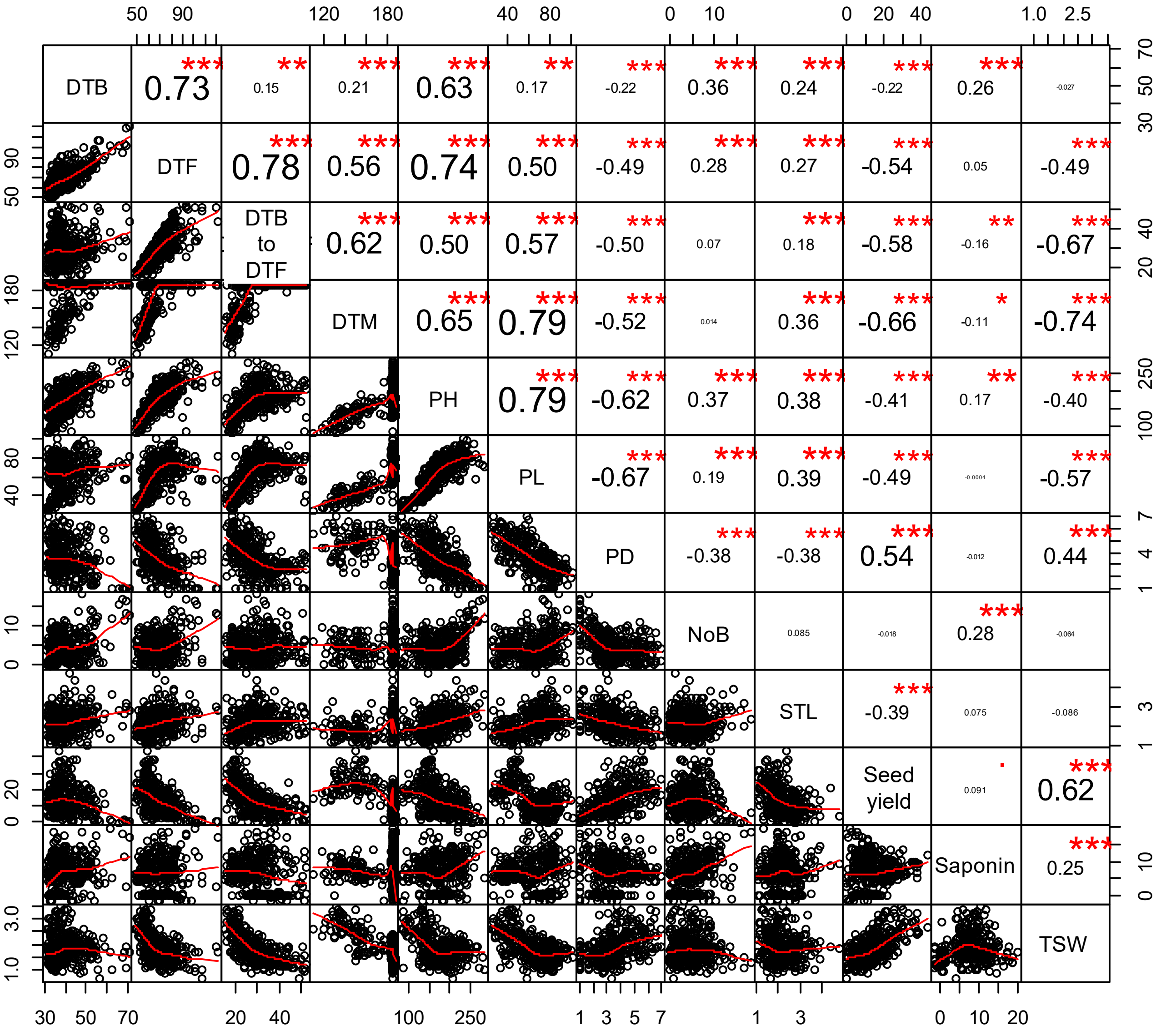


**Supplementary Fig. 10:** Pearson correlations among 12 quinoa traits. Best linear unbiased estimates across two years were used. Below the diagonal, scatter plots are shown with the fitted line in red. Above the diagonal, the Pearson correlation coefficients are shown with significance levels, *** =P<0.001, **=P<0.01.

(A)

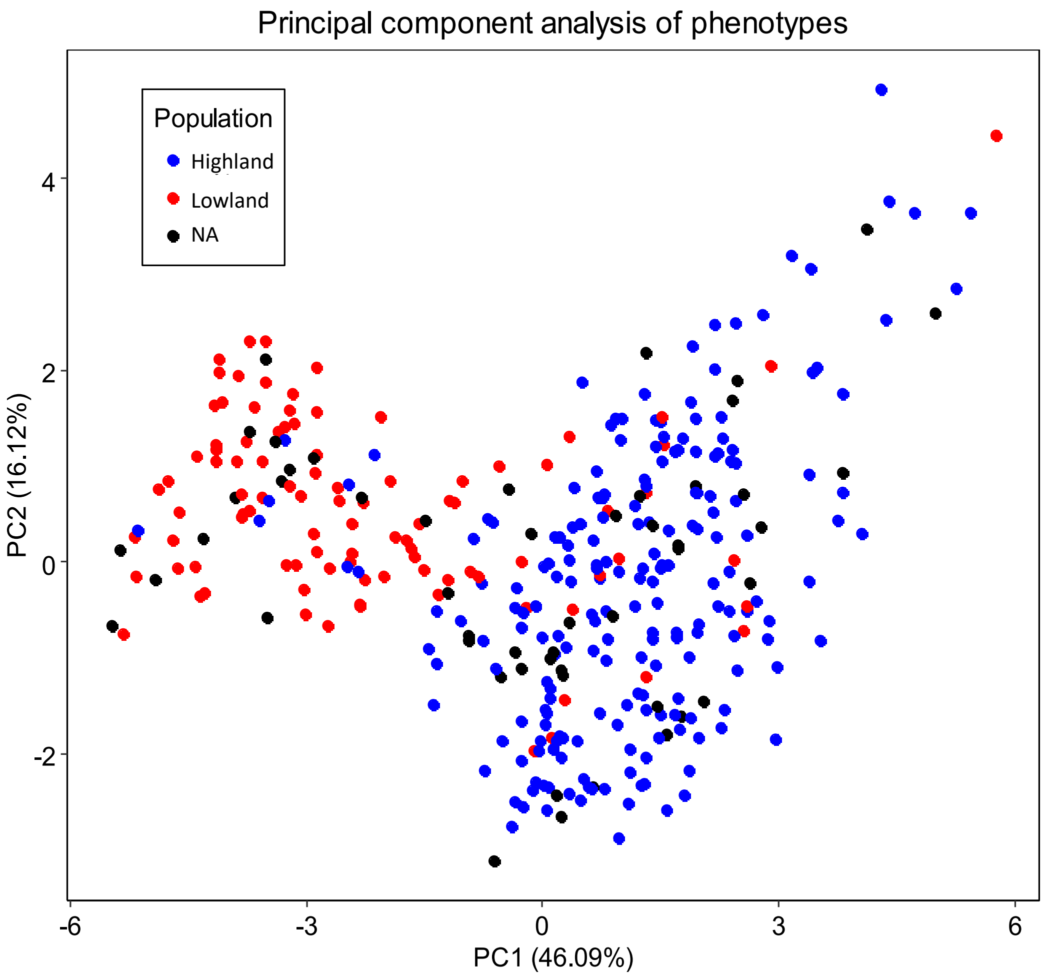


(B)

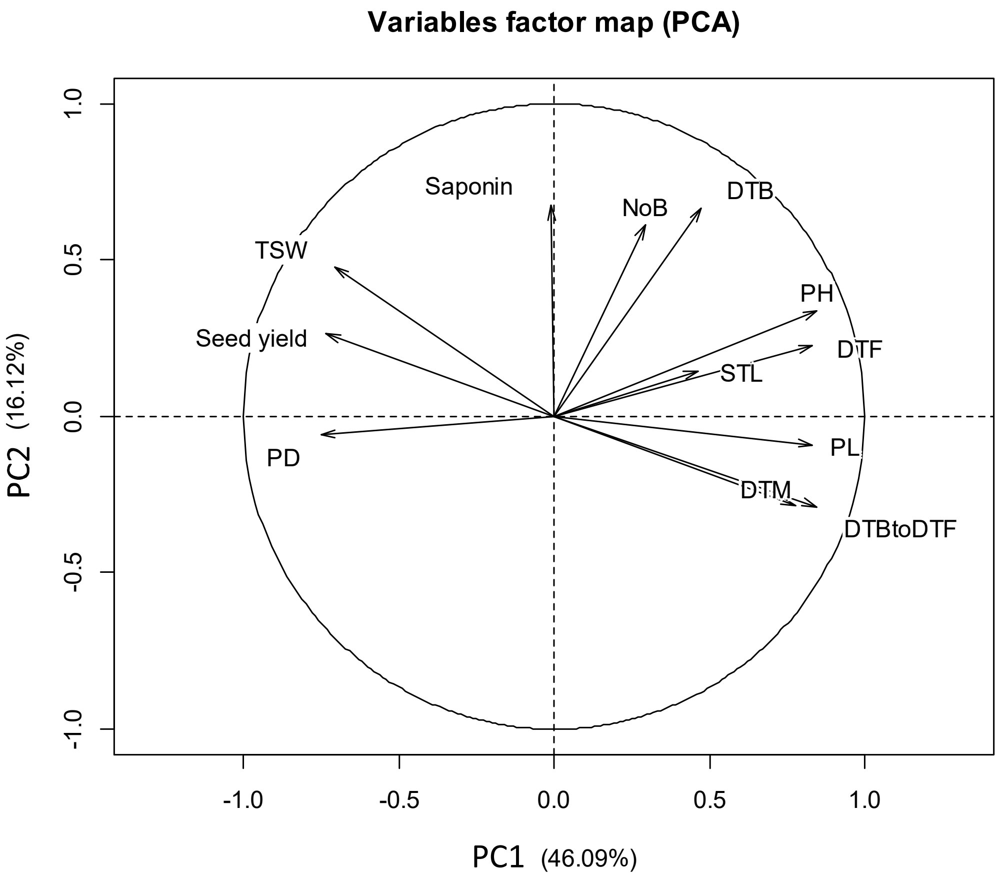


**Supplementary Fig. 11:** PCA of 12 quantitative phenotypes. A: Individual factor map colored according to populations identified from SNP analysis. B: Variables factor map of the PCA.

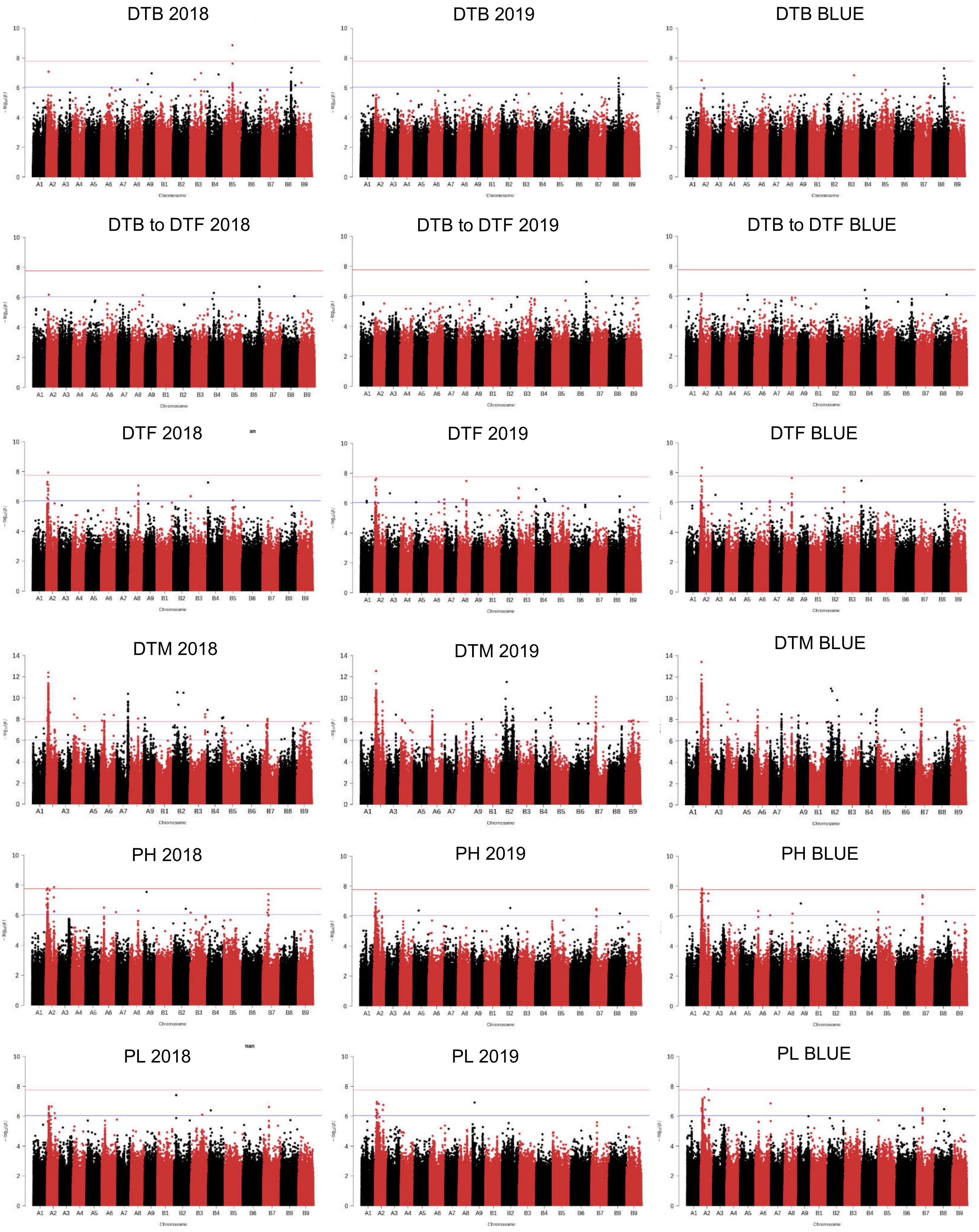


**Supplementary Fig. 12**: *cont.*

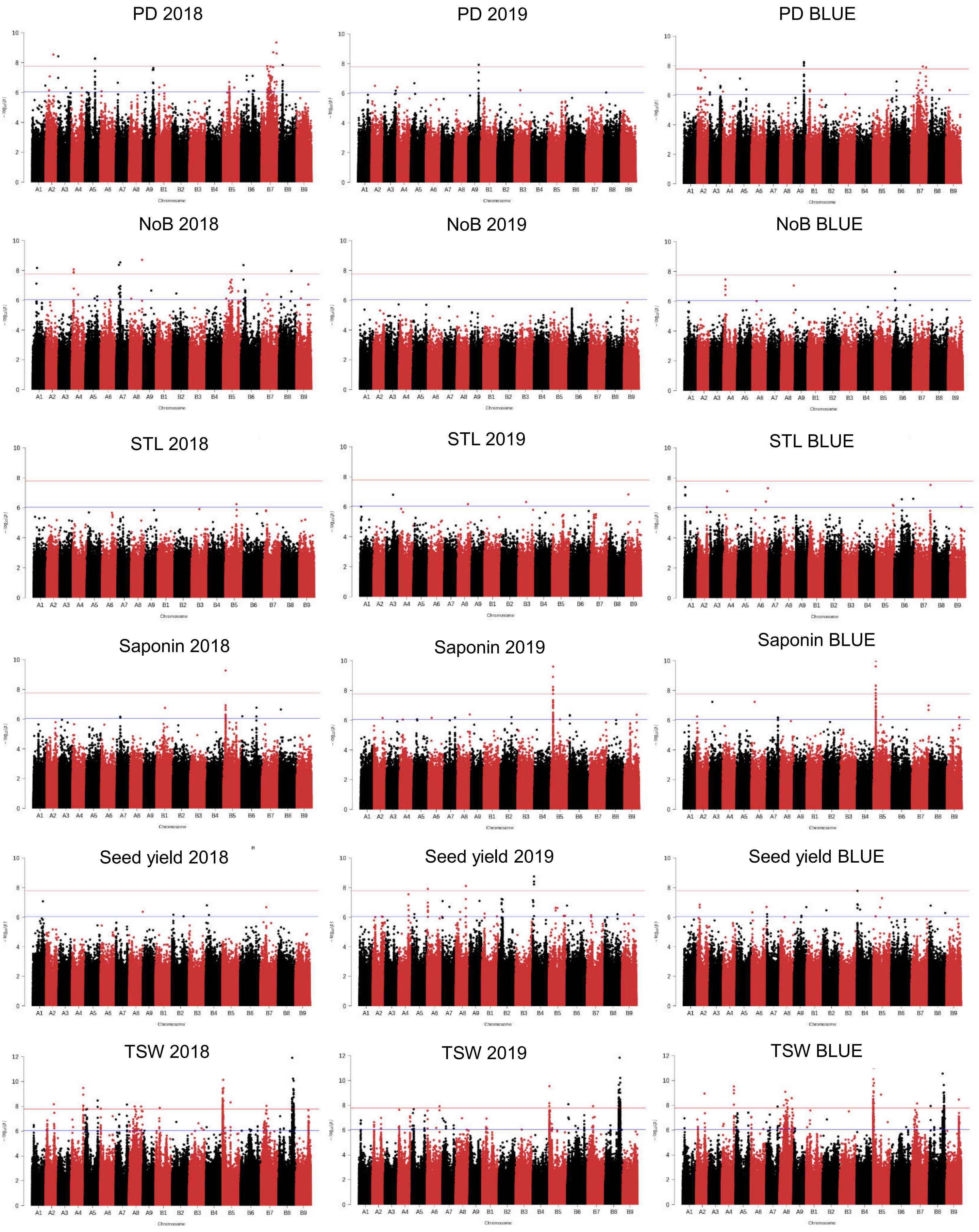


**Supplementary Fig. 12:** *cont.*

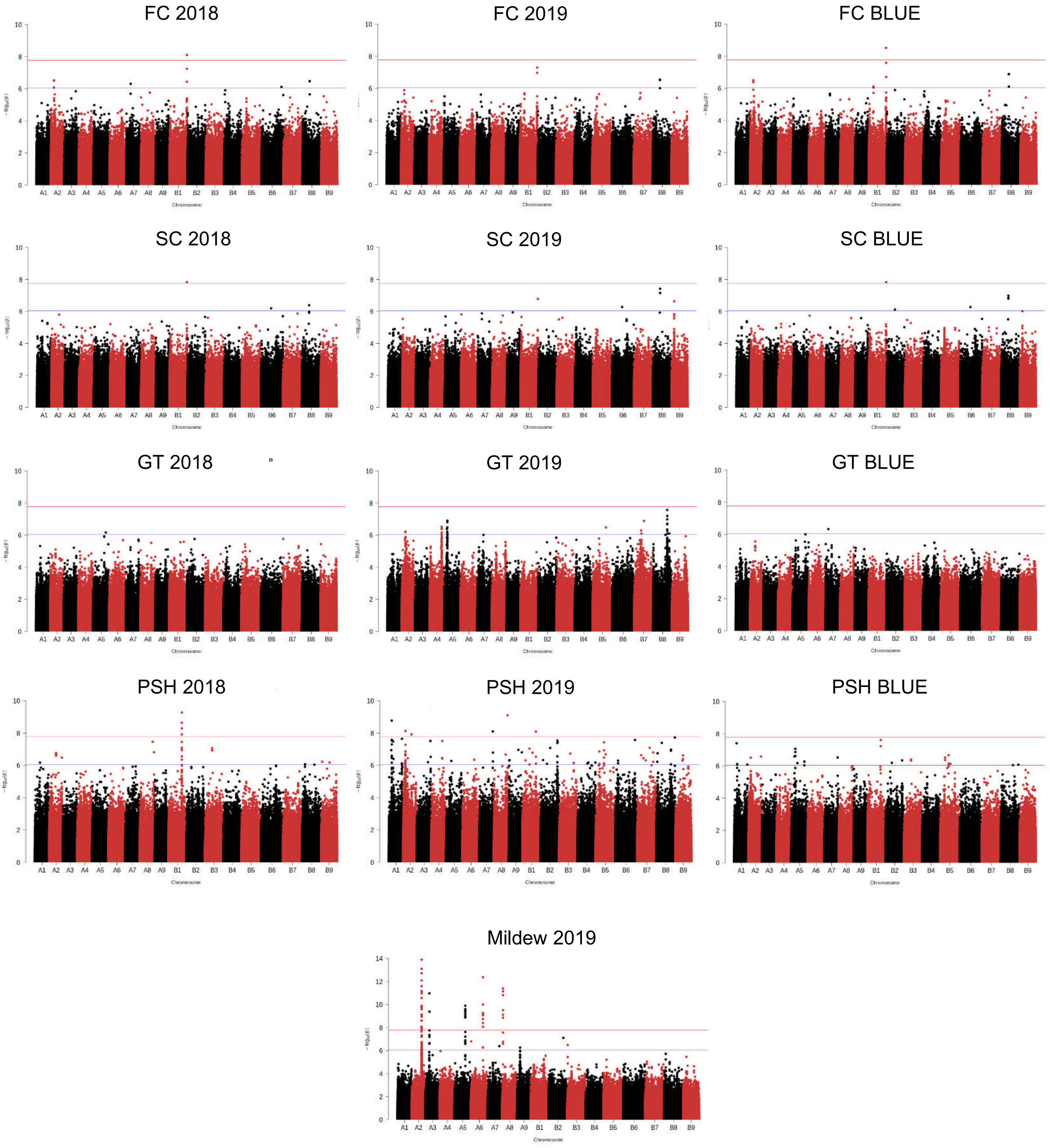


**Supplementary Fig.** 12:Manhattan plots from GWAS with data from 2018 (left), 2019 (center), and the mean of both years (right): The blue horizontal line indicates the suggestive threshold $-\log_{10}(8.98E-7)$. The red horizontal line indicates the significant threshold (Bonferroni correction) $-\log_{10}(1.67e-8)$.

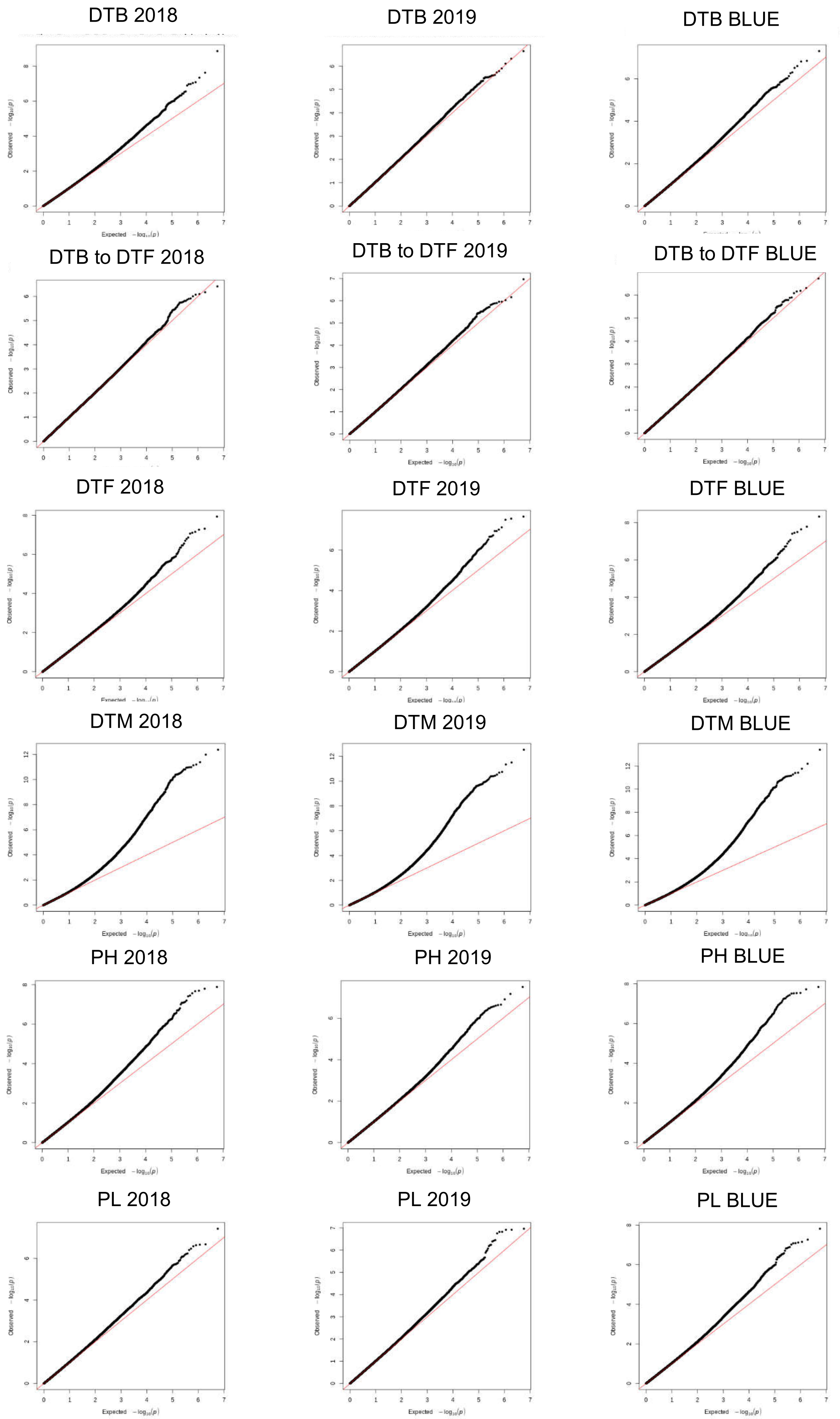


Supplementary Fig. 13: *cont*.

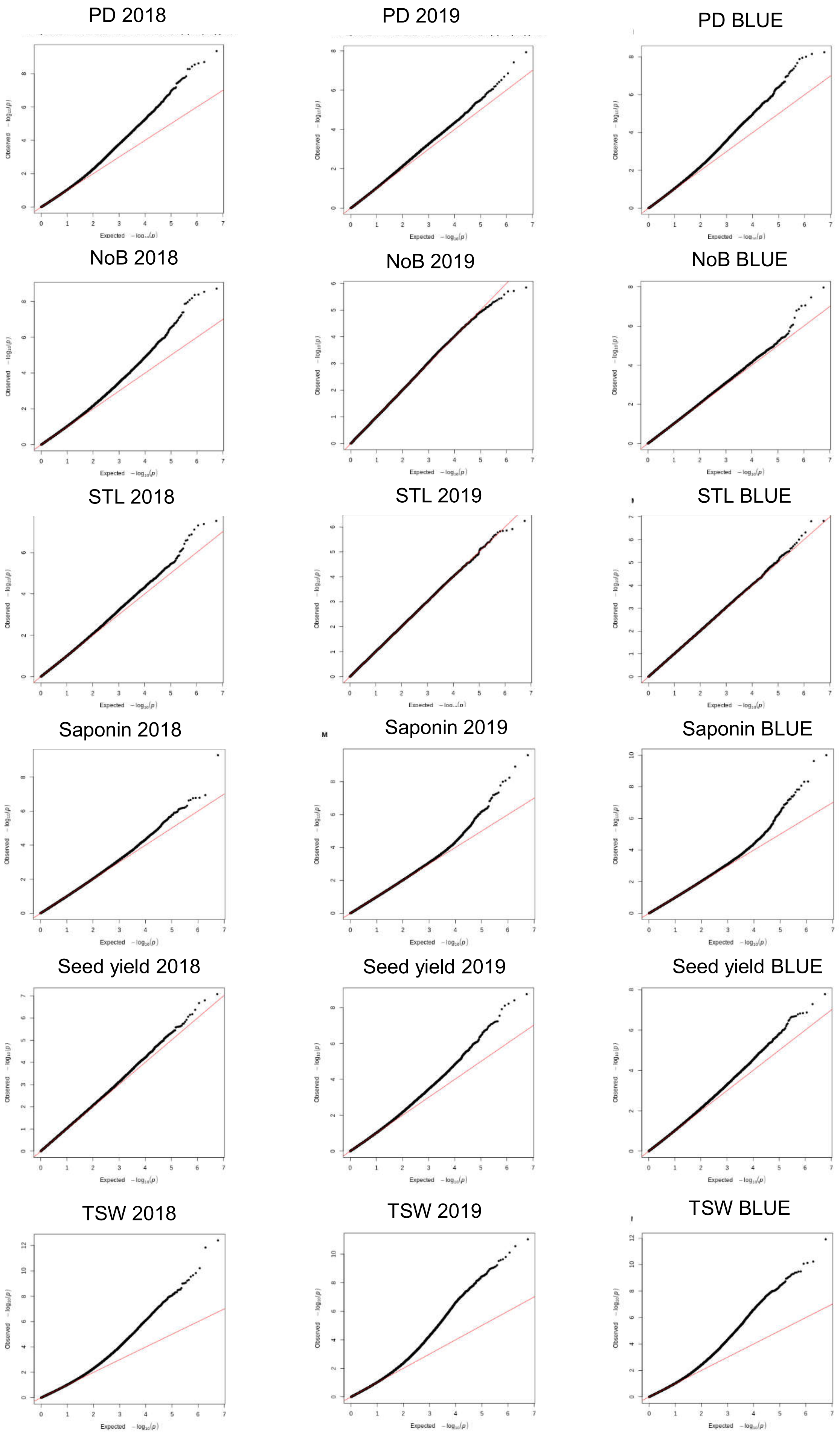


Supplementary Fig. 13: *cont.*

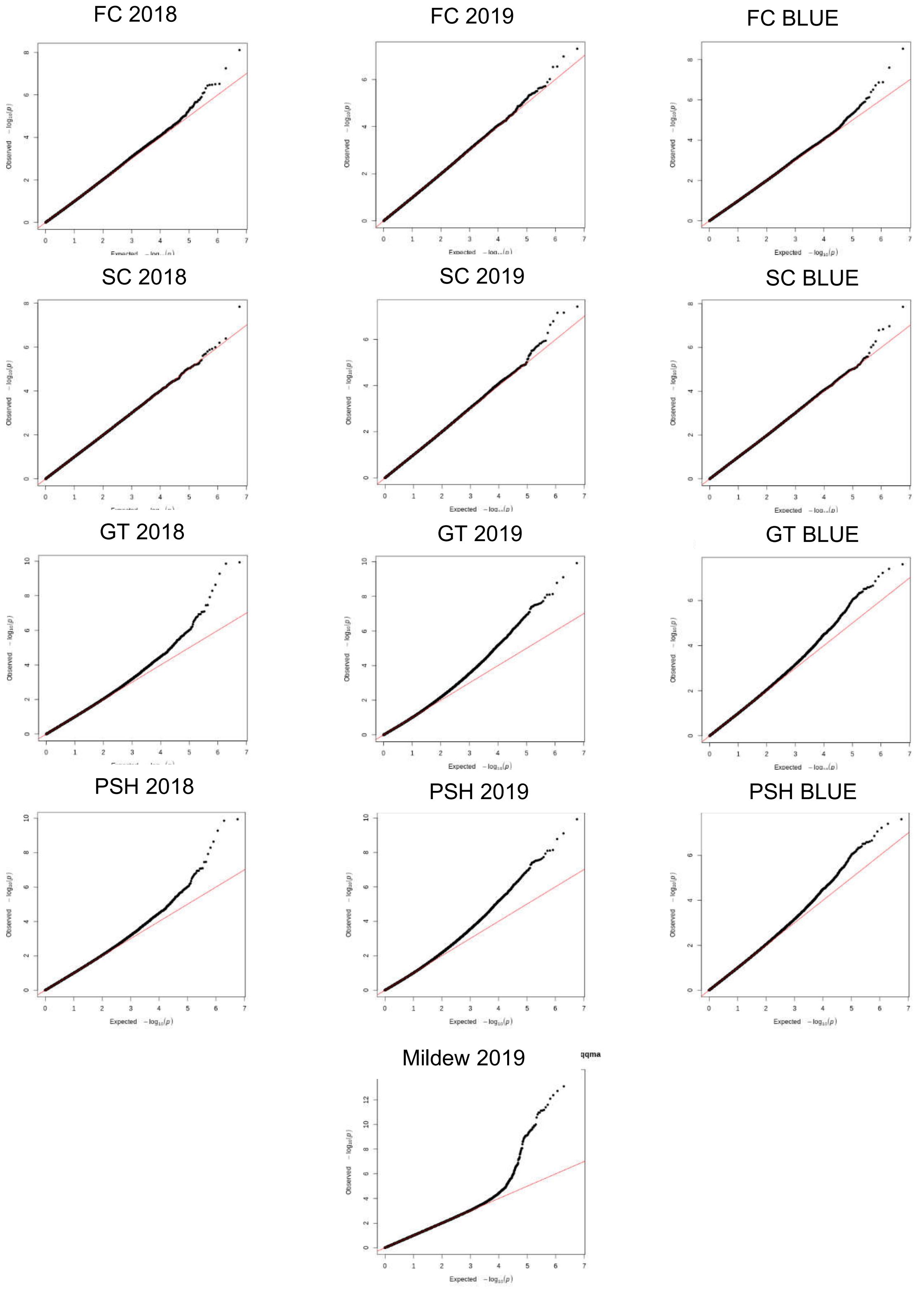


**Supplementary Fig. 13**: Quantile-quantile plots of GWAS in two years, 2018 (left) and 2019 (center), and BLUE (right).

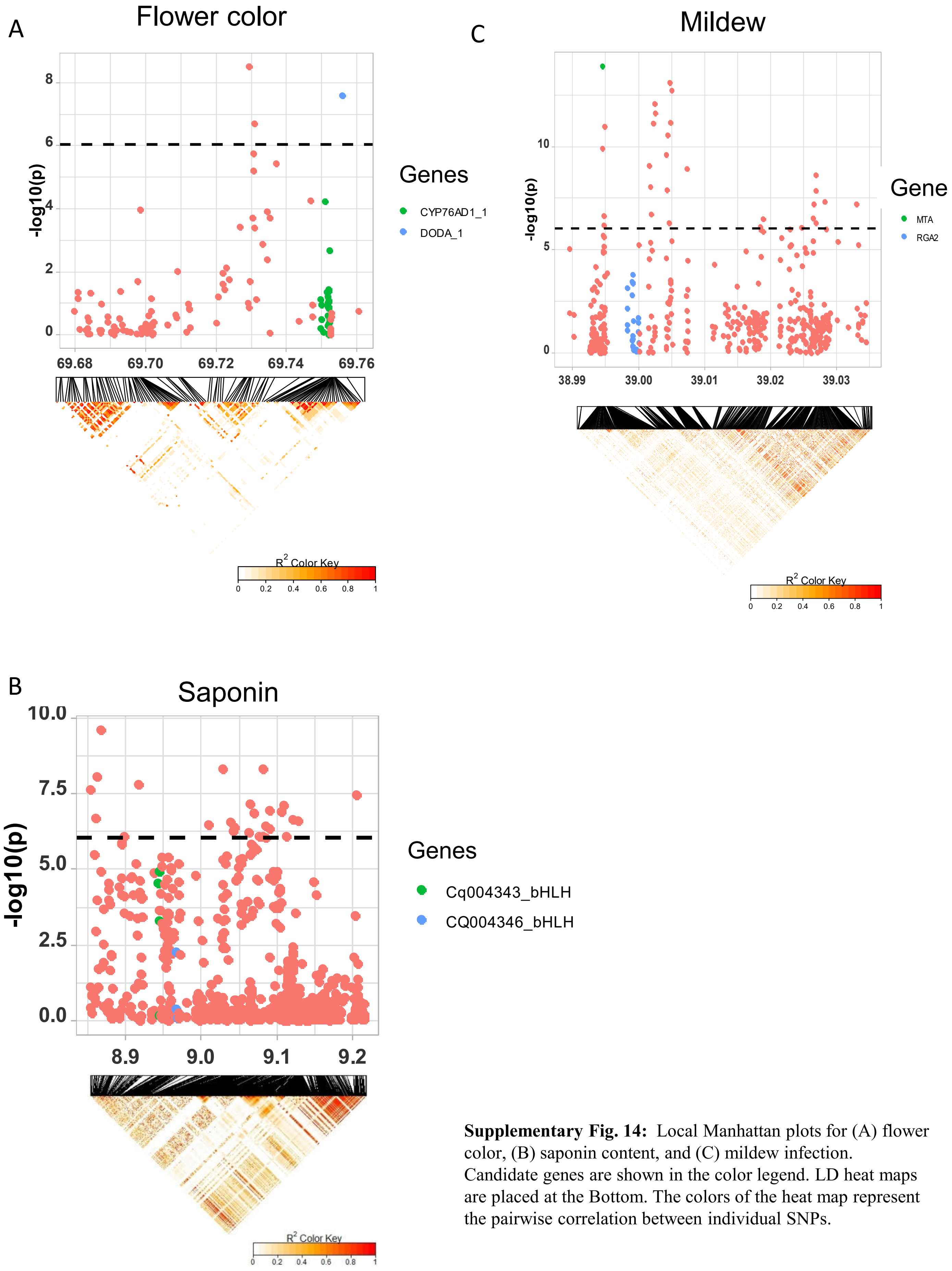


**Supplementary Fig. 14:** Local Manhattan plots for (A) flower color, (B) saponin content, and (C) mildew infection. Candidate genes are shown in the color legend. LD heat maps are placed at the Bottom. The colors of the heat map represent the pairwise correlation between individual SNPs.

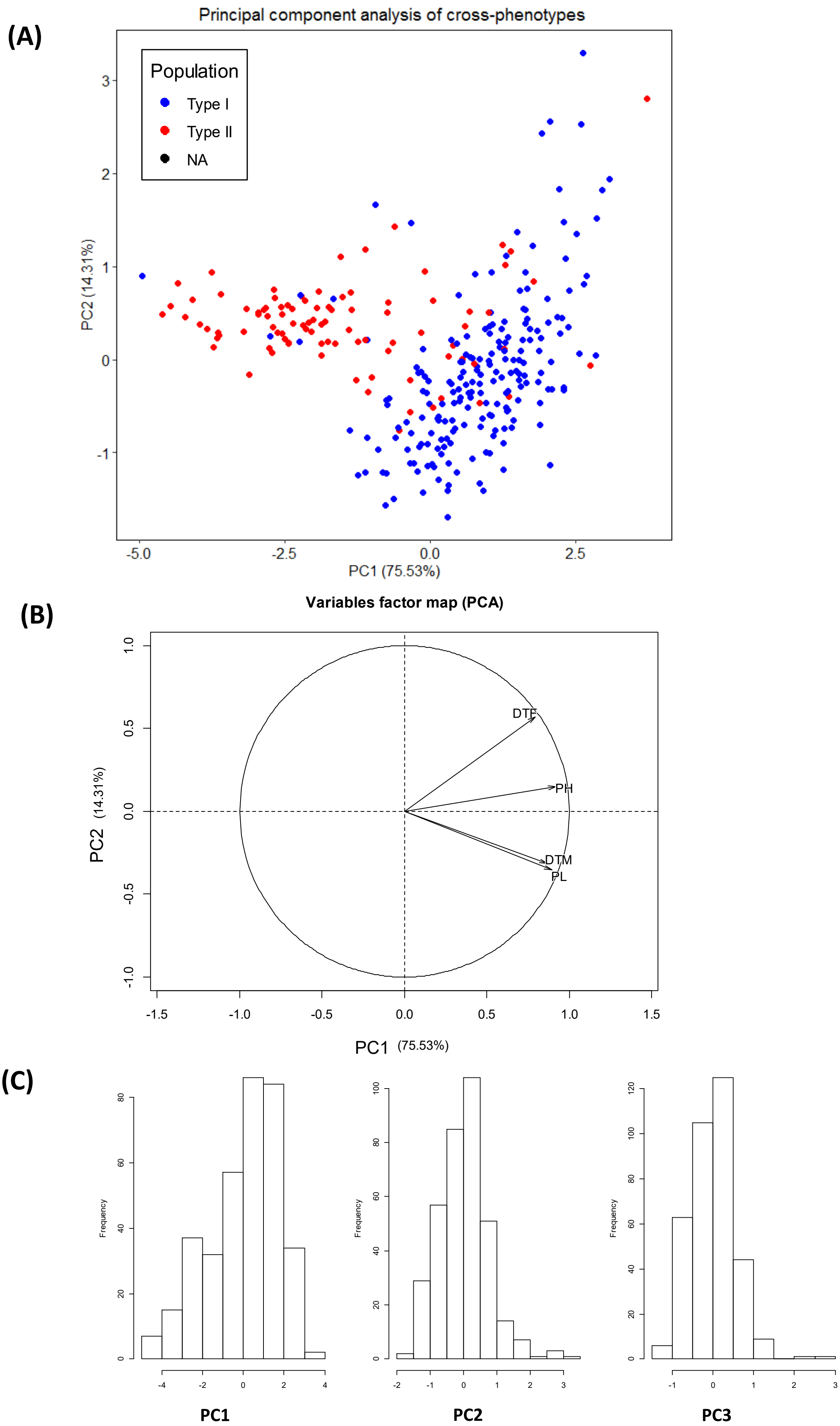


**Supplementary Fig. 15:** PCA of 4 quantitative traits (DTF, DTM, PH, and PL). A: Individual factor map, B: variables factor map of the PCA, C: distribution of the first three principal components which were used for GWAS analysis.

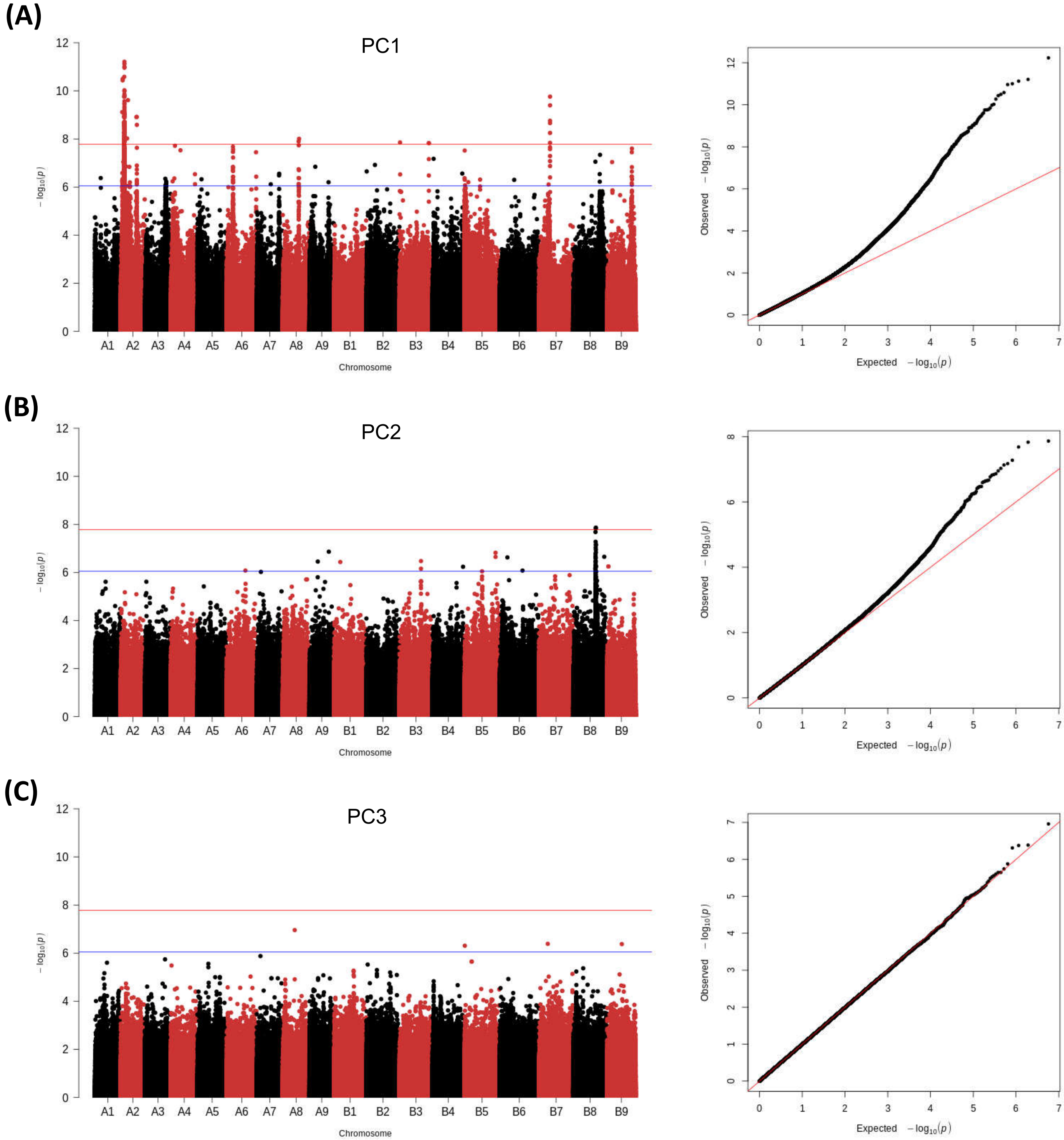


**Supplementary Fig. 16:** GWAS analysis of principal components, PC1 (A), PC2 (B), PC3 (C): Manhattan plots (left), and quantile-quantile plots (right): The blue horizontal line in the Manhattan plots indicates the suggestive threshold -log10(8.98E-7). The red horizontal line indicates the significance threshold (Bonferroni correction) -log10(1.67e-8).

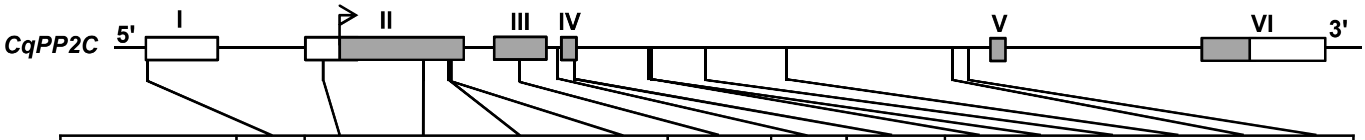


| UTR/Exon/Intron | Exon 1 | Exon 2 | | | | Exon 3 | Intron 3 | Exon 4 | Intron 4 | | | | | |
|---|---|---|---|---|---|---|---|---|---|---|---|---|---|---|
| SNP position | -1323 | -93 | 587 | 773 | 779 | 1252 | 1520 | 1637 | 2152 | 2160 | 2543 | 3101 | 4243 | 4346 |
| AA substitution | | | Arg196Arg | Ile258Met | Val260Val | Asp345Gly | | Ser440Ser | | | | | | |
| Hap1 | A | C | G | G | T | A | G | C | A | C | G | A | A | T |
| Hap2 | A | C | A | G | T | A | G | C | A | C | G | A | A | T |
| Hap3 | A | C | A | A | T | A | G | A | A | C | A | A | A | T |
| Hap4 | C | T | A | G | T | A | T | C | A | T | G | T | A | T |
| Hap5 | C | T | A | G | A | G | T | C | T | C | G | A | T | C |
| Missense variant | | | | * | * | * | | | | | | | | |

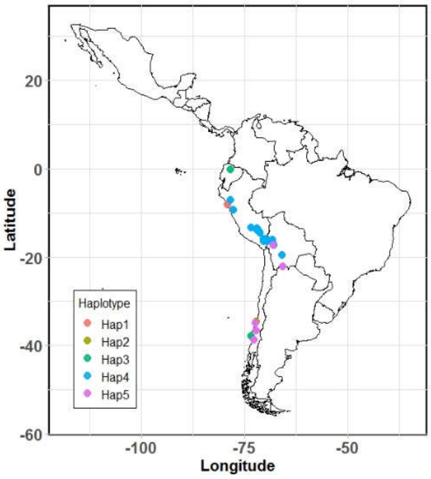


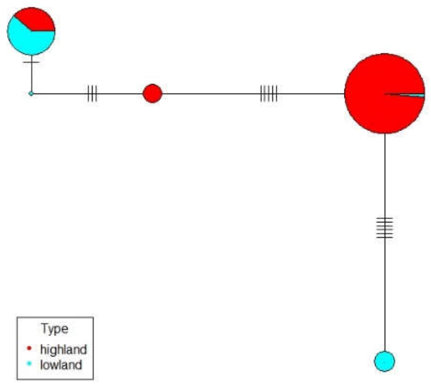


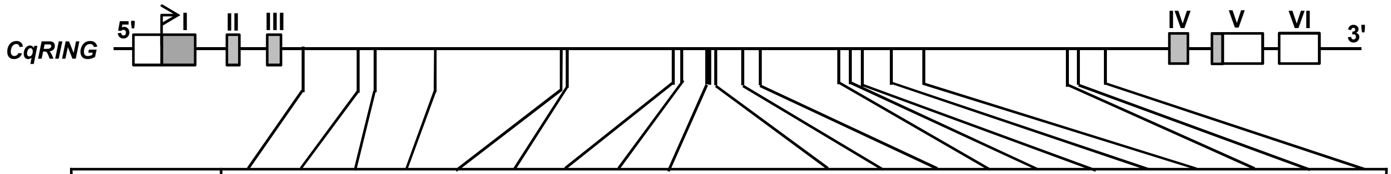


| UTR/Exon/Intron | Intron 3 | | | | | | | | | | | | | | | | | | | | |
|---|---|---|---|---|---|---|---|---|---|---|---|---|---|---|---|---|---|---|---|---|---|
| SNP position | 539 | 749 | 813 | 1045 | 1523 | 1548 | 1955 | 1980 | 2082 | 2084 | 2114 | 2221 | 2281 | 2586 | 2625 | 2670 | 2772 | 2782 | 2908 | 3455 | 3499 | 3593 |
| AA substitution | | | | | | | | | | | | | | | | | | | | | | |
| Hap1 | C | A | T | T | C | A | T | C | T | G | T | T | A | G | C | T | G | A | G | A | T | C |
| Hap2 | C | A | T | T | C | A | A | C | T | G | T | T | A | G | C | T | G | A | G | A | T | C |
| Hap3 | C | A | T | T | C | A | T | C | T | G | T | T | A | A | C | T | G | A | G | A | T | C |
| Hap4 | C | G | T | T | C | G | A | C | T | A | C | T | A | G | C | A | G | A | G | A | T | C |
| Hap5 | C | G | T | T | C | G | A | T | T | G | T | T | A | G | C | A | G | A | A | A | C | C |
| Hap6 | G | G | A | T | C | G | A | C | T | G | T | T | T | G | T | A | G | A | G | A | T | T |
| Hap7 | G | G | A | C | T | G | A | C | A | G | T | G | A | G | C | A | A | G | G | G | T | T |

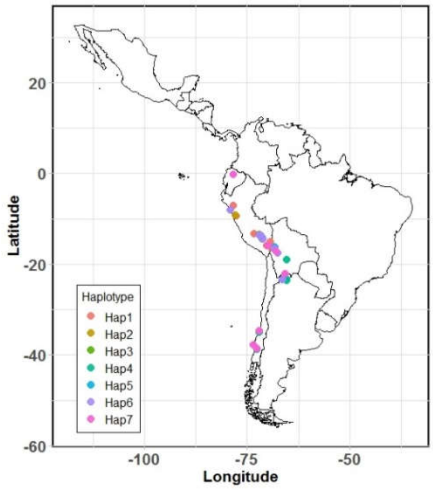


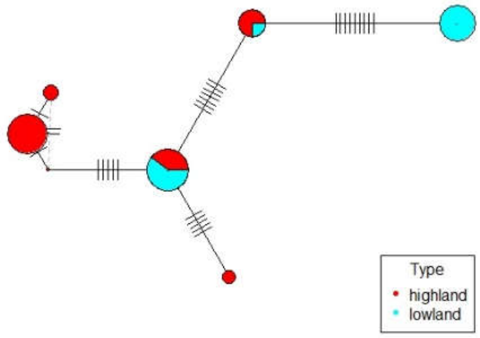


**Supplementary Fig. 17:** Haplotypes of two genes, *CqPP2C* and *CqRING* controlling seed size in quinoa. Geographic origin of the accessions and haplotype networks are displayed below the gene structure.
